# A systematic data-driven approach to analyze sensor-level EEG connectivity: Identifying robust phase-synchronized network components using surface Laplacian with spectral-spatial PCA

**DOI:** 10.1101/2021.08.10.455879

**Authors:** Ezra E. Smith, Tarik S. Bel-Bahar, Jürgen Kayser

## Abstract

Although conventional averaging across predefined frequency bands reduces the complexity of EEG functional connectivity (FC), it obscures the identification of resting-state brain networks (RSN) and impedes accurate estimation of FC reliability. Extending prior work, we combined scalp current source density (CSD; spherical spline surface Laplacian) and spectral-spatial PCA to identify FC components. Phase-based FC was estimated via debiased weighted phase-locking index from CSD-transformed resting EEGs (71 sensors, 8 min, eyes open/closed, 35 healthy adults, 1-week retest). Spectral PCA extracted 6 robust alpha and theta factors (86.6% variance). Subsequent spatial PCA for each spectral factor revealed seven robust regionally-focused (posterior, central, frontal) and long-range (posterior-anterior) alpha components (peaks at 8, 10 and 13 Hz) and a midfrontal theta (6 Hz) component, accounting for 37.0% of FC variance. These spatial FC components were consistent with well-known networks (e.g., default mode, visual, sensorimotor), and four were sensitive to eyes open/closed conditions. Most FC components had good-to-excellent internal consistency (odd/even epochs, eyes open/closed) and test-retest reliability (ICCs ≥ .8). Moreover, the FC component structure was generally present in subsamples (session × odd/even epoch, or smaller subgroups [n = 7-10]), as indicated by similarity of factor loadings across PCA solutions. Apart from systematically reducing FC dimensionality, our approach avoids arbitrary thresholds and allows quantification of meaningful and reliable network components that may prove to be of high relevance for basic and clinical research applications.

## 1. Introduction

Functional connectivity (FC) analyses seek to identify coordinated activity between spatially separated brain regions and, as such, can elucidate neural mechanisms that contribute to cognitive and affective processing in health and disease (Pizzagalli, 2011; Williams, 2017). There has been particular interest in characterizing stable patterns of interregional FC that emerge spontaneously during wakeful rest (Biswal, Zerrin Yetkin, Haughton, & Hyde, 1995; Raichle, 2015; Raichle et al., 2001). Identifying stable patterns of interregional FC has been accelerated by the use of multivariate techniques that can summarize thousands of functional connections into a few reliable components presumed to reflect latent functional networks. In resting participants, these networks have been referred to as resting state networks (RSNs; Lee, Smyser, & Shimony, 2013; Yeo et al., 2011).

Although functional magnetic resonance imaging (fMRI) has been the most frequently used technique for assessing RSNs, more recent studies have examined RSNs with electrophysiological methods including electroencephalogram (EEG) and magnetoencephalogram (MEG). Notably, recent work has suggested that M/EEG recordings detect somewhat different FC phenomena than neurometabolic assays (Forsyth et al., 2020; Mantini, Perrucci, Del Gratta, Romani, & Corbetta, 2007; Rizkallah, Amoud, Fraschini, Wendling, & Hassan, 2020; Coquelet et al., 2020; Marquetand et al., 2019). In particular, fMRI-based resting-state FC demonstrates the greatest convergence with electrophysiology at frequencies below (Hiltunen et al., 2014; Pan, Thompson, Magnuson, Jaeger, & Keilholz, 2013) and above (Fox, Foster, Kucyi, Daitch, & Parvizi, 2018; Kucyi et al., 2018) those EEG frequencies most prominent during wakeful rest (e.g., between 3 and 16 Hz; Congedo, John, De Ridder, & Prichep, 2010). This is not entirely surprising given that neurometabolic imaging and electrophysiology techniques measure neural activity at different temporal and spatial scales. Moreover, it is becoming increasingly apparent that high-level cognitive and motivational processes relevant to psychopathology are coded in the frequency and phase of coordinated neural activity between distal brain regions at sub-second time scales (Rouhinen et al., 2020; Ulhaas, Liddle, Linden, & Nobre, 2017). Unfortunately, most EEG FC studies have only examined a few hypothesized FC links from the thousands of possible FC links, or have focused on the sum of FC across all links, with very few systematically investigating the entirety of FC patterns inherent in high-dimensional EEG data. Moreover, most work to date has considered event-related rather than resting-state paradigms, and a comprehensive analysis of phase-synchronized FC during rest is lacking, especially when compared to the broad efforts in identifying fMRI-based RSNs (Mansour, Tian, Yeo, Cropsley, & Zalesky, 2021). Given the recent promising developments in using M/EEG connectivity biomarkers to predict clinical trajectories (e.g., Corlier et al., 2019; Whitton et al., 2018, 2019), a systematic identification of robust phase-based networks is urgently needed for streamlining biomarker development towards indices with psychometric validity and reliability adequate for clinical application.

### 1.1 Measuring functional connectivity in the human EEG

Although a comprehensive overview of connectivity metrics and methodology is beyond the scope of the present report, several excellent recent reviews exist (Bastos & Schoffelen, 2015; Marzetti et al., 2019; Siems & Siegel, 2020). In brief, there are four dominant approaches to measuring FC in electrophysiological recordings: 1) bivariate cross-spectra (e.g., Walter, 1963); 2) statistical dependency (e.g., autoregression) between broadband time series (e.g., Bressler & Seth, 2011); 3) correlation between temporally-filtered spectral amplitude envelopes (e.g., Hipp, Hawellek, Corbetta, Siegel, & Engel, 2012); and 4) phase clustering between electrode pairs (e.g., Lachaux, Rodriguez, Martinerie, & Varela, 1999). In general, no one connectivity metric outperforms all others, and the appropriateness of an FC metric for a given study depends on researcher hypotheses (Cohen, 2014; David, Cosmelli, & Friston, 2004; Wendling, Ansari-Asl, Bartolomei, & Senhadji, 2009). Whereas bivariate cross-spectra and phase clustering quantify synchronization of phase angles between two time series (i.e., phase-based FC), phase relationships are not specifically accounted for in most iterations of the other two approaches. This is notable since phase-based FC is unique to electrophysiological recordings, and is a primary hypothesized mechanism of neural communication (e.g., “communication through coherence”; Fries, 2015; Varela, Lachaux, Rodriguez, & Martinerie, 2001; Womelsdorf et al., 2007).

One promising phase-based FC metric is debiased-weighted Phase-Lag Index (dwPLI) that was introduced by Vinck and colleagues (2011). The dwPLI metric mitigates contributions of spurious FC from volume conduction and common generators (Bastos & Schoffelen, 2015; Nolte et al., 2004; Stam, Nolte, & Daffertshofer, 2007). Specifically, dwPLI accounts for the magnitude of phase angle differences based on their distance from the real polar axis, with phase clustering farther from the real axis being weighed more heavily, and by using a small offset to debias wPLI thereby countering the ‘bias’ effect resulting from a limited number of trials. Because of its resistance to quantifying spurious FC resulting from instantaneous volume conduction, dwPLI has become an increasingly popular phase-based FC metric (e.g., Alagapan et. al., 2019; González-Villar, Pidal-Miranda, Arias, Rodríguez-Salgado, & Carrillo-de-la-Peña, 2017; Portoles, Borst, & Van Vugt, 2018; van Driel, Gunsei, Meeter, & Olivers, 2017). Moreover, recent work has shown that dwPLI between EEG scalp sensors evidences adequate retest reliability (Hardmeier et al., 2014). Thus, for the current report, we chose dwPLI to measure FC in resting EEG recordings.

Reference-free EEG (Fein, Raz, Brown, & Merrin, 1988) has also been recognized as a technique for mitigating spurious FC arising from volume conduction and a common reference (e.g., Bastos & Schoffelen, 2015). Critical concerns about these problems were expressed decades ago (Essl & Rappelsberger, 1998; French & Beaumont, 1984; Leocani & Comi, 1999) and considered a ubiquitous confound, even when an average reference is used (Fein et al., 1988). Unsurprisingly, changing the EEG reference alters the observed FC patterns (Mehrkanoon, Breakspear, & Boonstra, 2014; Stam et al., 2007), although the true underlying neuronal generator patterns are obviously unchanged (e.g., Kayser & Tenke, 2010). The local Hjorth Laplacian (Hjorth, 1975), a computational variant of a surface Laplacian (second spatial derivative), produces sharper topographies indicating radially-directed current flow at scalp (current source density [CSD]; e.g., Kayser & Tenke, 2015b; Nunez et al., 1997; Perrin, Pernier, Bertrand, & Echallier, 1989; Tenke & Kayser, 2012). Scalp-based CSDs include all processes required for a complete description of neuronal current generators underlying the scalp EEG. Conversely, inverse source localization models, which are also independent of the EEG reference but notably dependent on biophysical model assumptions, can be precise but potentially incomplete or erroneous (Mahjoory et al., 2017; Tenke & Kayser, 2012). The scalp surface Laplacian (CSD) has also demonstrated critical advantages over surface potentials (i.e., referenced EEG) with regard to quantification of EEG phase (Tenke & Kayser, 2015). For these reasons, we employed a generic CSD transform using spherical splines as an initial processing step to obtain reference-free measures of EEG FC (Kayser & Tenke, 2015b; Tenke & Kayser, 2012, 2015).

### 1.2 Alpha and theta band networks

Only a handful of studies have specifically examined phase-based FC during wakeful rest, although this information is unique to M/EEG recordings and holds potential for clinical utility (Whitton et al., 2018, 2019). Of special interest, alpha (8 ~ 13 Hz) and theta (4 ~ 8 Hz) activity is frequently linked to neuropsychiatric syndromes and RSNs (Cavanagh & Frank, 2014; Cohen, 2011; Fries, 2015; Mantini et al., 2007; Sadaghiani & Kleinschmidt, 2016; Sadaghiani et al., 2010; Scheeringa et al., 2008).

Convergent evidence points to strong alpha band FC within occipital/visual regions, and between frontal and occipital regions (for a review, see Sadaghiani & Kleinschmidt, 2016). For alpha, Stam et al. (2007) found strong FC between occipital-parietal sensors at rest in MEG recordings for both phase-locking index (PLI) and imaginary coherency (iCOH). Using beamforming to source localize resting MEG, Hillebrand et al. (2012) found strong local PLI for 8-13 Hz alpha within medial occipital regions. A follow-up study (Hillebrand et al., 2016) analyzed FC for low (8 – 10 Hz) and high (10 – 13 Hz) alpha bands separately, with both frequency bands revealing a similar pattern of local FC near the cuneus, and long-range FC between the cuneus and medial prefrontal cortex (mPFC; Hillebrand et al., 2016). In contrast, Hardmeier et al. (2014) observed two distinct FC patterns for the low (8 – 10 Hz) and high (10 – 13 Hz) alpha bands during rest using dwPLI applied to average-referenced 256-sensor EEGs. Whereas low alpha was characterized by strong long-range FC between midline sensors over frontal, central, and occipital regions, high alpha was characterized by strong local FC amongst a cluster of medial parietal-occipital sensors, suggesting frequency-specific connectivity within the traditional alpha band. By comparison, theta band FC in resting EEG was pronounced locally in midfrontal brain regions (Hardmeier et al., 2014; Hillebrand et al., 2016; Vidaurre et al., 2018), with one MEG study failing to identify significant theta PLI in a small sample (*N* = 13; Hillebrand, Barnes, Bosboom, Berendse, & Stam, 2012).

Notably, these studies relied on averaging across fixed canonical frequency bands rather than empirically determining FC spectral bins. Unfortunately, this approach can obscure frequency-specific FC patterns if a priori frequency bands do not align with natural FC spectra. This is especially problematic if multiple RSNs operate at neighboring but nonetheless distinct frequencies, for instance, within narrow alpha and/or theta bands. On the other hand, multivariate approaches can identify natural (i.e., data-driven) FC frequencies, and summarize FC patterns into a connectivity component, interpreted here to reflect a RSN. In fact, the few studies analyzing M/EEG FC with multivariate approaches replicated the findings reported above, and identified additional candidate RSNs. For example, Mehrkanoon et al. (2014) identified seven RSNs in the resting EEG using a PCA variant. Two components overlapped with the occipital and midline alpha FC patterns identified by Hillebrand et al. (2016), with evidence for a spectral peak in the alpha band (10~12 Hz). An additional alpha component was identified with a peak at 12~13 Hz and an FC pattern consistent with the sensorimotor mu network (Pfurtscheller, Neuper, Andrew, & Edlinger, 1997). Mehrkanoon et al. (2014) also found a component with a midfrontal spatial pattern consistent with one previously seen for theta FC, however, it had spectral peaks in both the theta *and* alpha range. More recently, Vidaurre et al. (2018) used Hidden Markov Modeling (HMM) to identify RSNs with source-reconstructed MEG. One RSN demonstrated strong local FC in medial and lateral parietal regions at ~ 9 Hz, and another RSN was characterized by local FC within occipital regions at ~ 11 Hz, congruent with RSNs found by Mehrkanoon et al. (2014) and Hillebrand et al. (2016). Vidaurre et al. (2018) also identified a ~12 Hz sensorimotor component similar to that of Mehrkanoon et al. (2014), and a midfrontal-frontotemporal connectivity component spanning the delta and theta range (1~7 Hz).

Altogether, these findings converge on four spectral-spatial FC patterns characterized by: 1) a spectral peak at 10~11 Hz prominent over occipital regions; 2) a peak at 8~9 Hz spanning midline regions; 3) a ~12 Hz peak with central/sensorimotor regions; and 4) a delta-theta peak (2~8 Hz) within midfrontal regions. The few studies analyzing M/EEG FC with multivariate approaches found FC patterns that were grossly congruent with univariate approaches; however, multivariate techniques also identified additional RSNs not previously recognized.

### 1.3 Stability and reliability of EEG FC

Only recently have investigators began to examine the test-retest reliability of EEG FC at rest. For simplicity and interpretability, a high-dimensional connectivity dataset has to be reduced into a manageable number of test “items,” often a mean. In the M/EEG FC literature, for example, many reports have calculated a grand mean across all FC edges (i.e., “global” reliability). Several EEG studies have shown adequate test-retest reliability for global FC in the alpha band across one-week to two-year intervals, whereas global theta reliability has been less consistent^1^ (Dunkin, Leuchter, Newton, & Cook, 1994; Hardmeier et al., 2014; Kuntzelman & Miskovic, 2017; Marquetand et al., 2019; van der Velde, Haartsen, & Kemner, 2019). By comparison, reliability of individual FC links was in the poor to questionable range in one study with infants (van der Velde et al., 2019). To the best of our knowledge, regional variability in FC has been ignored by analyses examining individual FC links and global FC. In fact, several studies have revealed that FC test-retest reliability depends on region (Hardmeier et al., 2014; Holler et al., 2017; Roberts, Fillmore, & Decker, 2016; van der Velde et al., 2019), with the strongest FC links also demonstrating greater reliability (Garces et al., 2016). By comparison, multivariate techniques automatically organize FC links into reliable components/RSNs, thereby obviating some of the shortcomings of univariate analyses that do not account for regional differences in FC reliability. Taken together, whereas initial findings on test-retest reliability of global FC at alpha and theta bands have been promising, the reliability and robustness of FC patterns (i.e., putative RSNs) has not been systematically examined.

### 1.4 Present study

This report focused on the systematic characterization of theta and alpha connectivity to interrogate the reliability of spatial FC patterns. To this end, a combination of CSD-transformation and PCA was used to identify and quantify connectivity components within thousands of individual FC links, a technique which we termed CSD-fcPCA. PCA is a linear multivariate method for identifying and summarizing common variance in electrophysiological data that has played an important role in consolidating the construct of a component (Chapman & McCrary, 1995; Donchin, 1966; Donchin & Heffley, 1978; Kayser & Tenke, 2003; Tenke & Kayser, 2005; Van Boxtel, 1998). PCA has successfully guided electrophysiological research in temporal, spectral, spatial, and temporal-spectral domains (Kayser & Tenke, 2006b, 2015c; Kayser, Tenke, & Debener, 2000; Tenke et al., 2008). In this study, we extended the PCA approach to identify frequency-specific connectivity components from sensor-level FC data. Cross-validation was performed on data subsamples to evaluate the consistency of PCA solutions (e.g., Barry, De Blasio, & Karamacoska, 2019) and to rule out components that may be unique to only a few participants (e.g., Smith et al., 2020). Components were also evaluated for internal consistency and test-retest reliability using Intraclass Correlations (ICCs; Shrout & Fleiss, 1979), and compared with regard to resting conditions (eyes open versus closed).

## Method

### 2.1 Participants

Details regarding participant recruitment and selection are presented in Tenke et al. (2017). Reported data are from 35 English-speaking healthy adults (21 female, aged 18 to 65 years [Mean ±SD, 35.6 ±14.3], 10 to 20 years of education [15.3 ±2.3]) of diverse ethnicity (23 White, 7 African-American, 3 Asian, 2 Other) and mixed handedness (5 left-handed; Edinburgh Handedness Inventory laterality quotient +66.6 ±53.8, range ‒100.0 to +100.0; Oldfield, 1971) who were enrolled in the NIMH-funded project *Establishing Moderators and Biosignatures of Antidepressant Response in Clinical Care* (EMBARC; Trivedi et al., 2016). Participants were locally recruited and tested at four data collection sites (Tenke et al., 2017): Columbia University Medical Center in New York (CU; *n* = 10), Massachusetts General Hospital in Boston (MG; *n* = 9), University of Texas Southwestern Medical Center in Dallas (TX; *n* = 9), and University of Michigan in Ann Arbor (UM; *n* = 7). All participants had no history of any DSM-IV disorder (First, Spitzer, Gibbon, & Williams, 1996). Each participant had two resting EEG recordings separated by about one week. Each EEG recording included two eyes-closed (C) and two eyes-open (O) blocks arranged in a fixed OCCO order, with each block lasting 2 min. Participants also completed additional biometric assays and self-report questionnaires that are not pertinent to the present report. The study was conducted in accordance with the Declaration of Helsinki, was approved by the institutional review board at each testing site, and all participants provided informed consent.

### 2.2 EEG processing and CSD transform

All EEG pre- and postprocessing steps, including unification of EEG montage and acquisition parameters, have been detailed in Tenke et al. (2017). Briefly, data from the different collection sites were converted to a common 71-channel EEG montage (10-5 system; Jurcak, Tsuzuki, & Dan, 2007; see Kayser & Tenke, 2015a, Fig. 1 minus the Nose reference site and P9/10 instead of P09/10) and visually inspected for recording artifacts. Missing, bad or bridged channels (Alschuler, Tenke, Bruder, & Kayser, 2014) were replaced by spherical spline interpolation (Perrin et al., 1989). The continuous EEG data were then blink-corrected via spatial singular value decomposition (NeuroScan, 2003) and segmented into 2-s epochs with 75% overlap. Epochs were band-passed at 1-60 Hz (24 dB/octave). A semi-automated reference-free approach identified isolated EEG channels containing amplifier drift, residual eye activity, muscle or movement-related artifacts on a trial-by-trial basis. Channels containing artifact were replaced by spline interpolation if less than 25% of all channels were flagged; otherwise, the epoch was rejected. An automatic threshold (±100 μV) applied to all EEG channels was used to exclude deviant trials. Lastly, an automated ICA-based correction was also applied that subtracts components suspicious for any additional non-neurogenic artifacts (i.e., EMG, bad electrodes) without removing additional data segments (Smith, Reznik, Stewart, & Allen, 2017).

**Figure 1.**
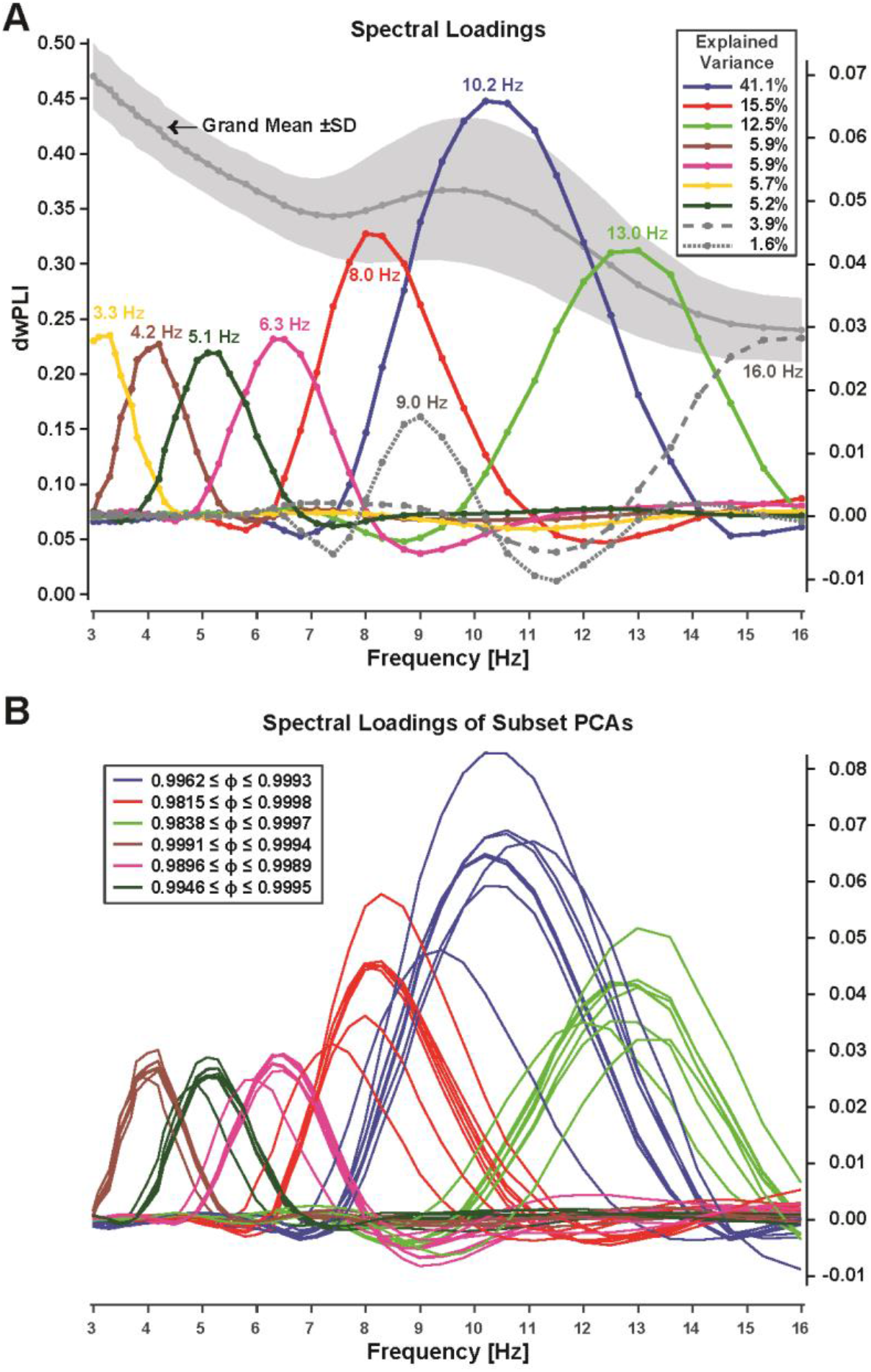
Spectral FC components identified via step-one PCA. **A.** Grand mean (±SD) debiased weighted phase-locking index (dwPLI) across all 2,485 edges (left ordinate) and spectral loadings of the first 9 components extracted (right ordinate). Inset shows explained variance. Loadings were identified by peak frequency (listed in same color next to waveforms). Line markers correspond to the logarithmically-spaced frequency bins. **B.** Consistency of step-one spectra FC components across data subsets (see text). Line color of subset loadings correspond to those matching one of the six overall PCA loadings (A) considered for further analysis. Inset shows ranges of Tucker congruence coefficients (ϕ) for each spectral loading considered, indicating equality.

To eliminate or substantially reduce the deleterious effects of volume conduction and common reference on phase-relationships between EEG recording sites (e.g., Bastos & Schoffelen, 2015; Nunez et al., 1997; Tenke & Kayser, 2012, 2015), a reference-free CSD transform (spherical spline surface Laplacian; Perrin et al., 1989) was applied to all artifact-free EEG epochs using the Matlab implementation of the CSD toolbox (Kayser, 2009; Kayser & Tenke, 2006b) with default settings (spline flexibility, *m* = 4; smoothing constant, λ = 10^−5^; Legendre polynomial, number of iterations = 50)^2^. CSDs are based on the second spatial derivative of surface potentials and represent the magnitude of radial current flow entering (source) and leaving (sink) the scalp, constituting a unique, unambiguous measure of neuronal activity at sensor level (e.g., Tenke & Kayser, 2012). In contrast to inverse source localization techniques, CSDs do not rely on a specific volume conductor model (Carvalhaes & de Barros, 2015) and are therefore a conservative means to faithfully characterize both shallow and deep neuronal generators underlying a scalp potential topography, including partially-closed fields (for a review, see Tenke & Kayser, 2012). We and others have repeatedly emphasized the benefits of CSD methods over using reference-dependent surface potentials (Burle et al., 2015; Cohen, 2015; Hagemann, Naumann, & Thayer, 2001; Kayser & Tenke, 2015c; Smith et al., 2017; Velo, Stewart, Hasler, Towers, & Allen, 2012).

### 2.3 Phase-synchronization and functional connectivity

A Morlet wavelet procedure (Cohen, 2014) was used to extract the analytic signals which are the basis for calculating debiased-weighted phase-lag index (dwPLI; Vinck et al., 2011), a measure of functional connectivity between brain regions. Larger dwPLI values, which range from 0 to 1, indicate consistent differences in phase lag between electrode pairs; however, dwPLI coefficients do not indicate connectivity direction. A total of 40 Morlet wavelets were logarithmically-spaced and ranged from 2 to 50 Hz. Wavelet cycles ranged from 3 to 10, with lower frequencies having fewer cycles. As the focus was on FC within theta and alpha bands, we restricted data analysis to a 3-16 Hz frequency range, resulting in 42 frequency bins^3^. A dwPLI value was calculated for each possible pair of 71 electrodes ([71 × 70]/2 = 2,485 unique pairs or edges) from a 1-s segment from the middle of each 2-s EEG epoch. The adjacent outer 0.5-s segments were excluded to mitigate edge artifacts.

### 2.4 Two-step connectivity PCA

PCA decomposes a two-dimensional EEG-FC data matrix into a set of loadings (along the variable dimension [columns]) and corresponding scores (along the case dimension [rows]) that provide a weight for each observation, which together constitute a set of FC components (see Kayser & Tenke, 2003, for temporal PCA decomposition that follows the same decomposition principle). A loadings vector (i.e., one for each extracted factor) reflects a linear covariance pattern in multivariate space. When using an unrestricted factor extraction and rotation (Kayser & Tenke, 2003), there are as many loadings as there are variables, provided the number of cases exceeds the number of variables and there are no linear dependencies among variables. For each loading vector, there are as many scores as there are observations, which reflect the strength of a given loading pattern for each observation.

Given the substantial number of variables per condition (i.e., 42 frequencies × 2,485 edges = 104,370 variables) compared to the number of cases (i.e., 35 participants × 2 eyes open/closed conditions × 2 testing sessions = 140 cases), a two-step approach was chosen so that the number of cases exceeded the number of variables (Gorsuch, 1983). First, FC data was decomposed into latent spectral FC components (i.e., analogous to a frequency PCA using power or amplitude spectra; Tenke & Kayser, 2005), entering only frequencies as variables whereas edges, conditions, sessions and participants were entered as cases. For the second PCA, raw FC data matrix was filtered^4^ by a latent spectral factor, and then rearranged so that FC links/edges were variables, and frequencies, conditions, sessions and participants were cases. This spectrally-filtered data matrix was then decomposed into latent connectivity components each with a unique spectral-spatial profile. In brief, the first PCA step was used to identify specific frequency-bands with high-variance dwPLI; these step-one spectral FC components were then back-projected back into dwPLI space, effectively filtering FC links by frequency, and a second PCA identified spatial FC components from all possible bivariate FC coefficients between sensor pairs.

The above considerations resulted in the following data matrices submitted to PCA at each step. For step one, variables consisted of 42 discrete frequency bins between 3 and 16 Hz to summarize spectral FC covariance across 695,800 cases (observations) stemming from participants (35), recording sessions (2), conditions (2), odd/even epochs (2), and all connectivity pairs (2485 edges). Prominent high-variance components from the step-one PCA were then back-projected into original data units (i.e., dwPLI)^5^, yielding a full data matrix that represented connectivity strength for a specific frequency. For step two, the back-projected, component-filtered data matrix was then reorganized by arranging all connectivity pairs as variables and all other dimensions (i.e., participants, sessions, conditions, odd/even epochs, frequency bins) as cases, resulting in a new matrix (11,760 cases × 2,485 variables) for factoring the spatial connectivity covariance (i.e., pairwise connectivity strength between all 71 scalp sensors for a specific frequency pattern). Therefore, components extracted during the step-two PCA represent frequency-specific spatial FC patterns, with corresponding scores reflecting a concise summary for the strength of a particular, frequency-specific network pattern for each case (i.e., participant, recording session, condition, and odd/even epoch). As with other PCA solutions, these component scores can then be easily subjected to follow-up statistics (e.g., Kayser & Tenke, 2003; Kayser et al., 2014).

In addition to the PCA solution calculated over the entire dataset, subsets of data were also submitted to PCA to evaluate consistency of component loadings. Specifically, the two-step PCA procedure was repeated for individual data collection sites and recording sessions divided by odd/even trials. These eight additional subset PCAs (4 collection sites, 4 session × epoch [odd/even trials]) were calculated using the average FC across all resting epochs available. There were a minimum of 74 trials available for each condition in session-one recordings, and 44 trials available for each condition in session-two recordings. After obtaining the step-one PCA solutions for each subset for comparison, step-two PCAs conducted for data subsets were filtered by the spectral FC components of the step-one PCA solution derived from the group-level analysis (i.e., from the entire dataset). In this way, slight variations in step-one solutions for different subsets of data were avoided so that the stability of step-two solutions could be evaluated without introducing confounds stemming from different step-one solutions.

To be consistent with our prior CSD-PCA approaches for temporal, spectral, spatial and time-frequency EEG data (Kayser & Tenke, 2005, 2006b; Kayser et al., 2014; Tenke et al., 2011) the main group-level PCA was based on an unrestricted solution followed by Varimax rotation (Kaiser normalization) of covariance loadings (Tenke & Kayser, 2005). Component extraction was restricted to 50 for step-two PCAs calculated on data subsets to reduce computation time after verifying that only trivial differences were observed between an unrestricted CSD-fcPCA and a CSD-fcPCA restricted to 50 components, given that the vast majority of extracted factors of an unrestricted PCA solution will explain little and most likely not theoretically-meaningful variance (Kayser & Tenke, 2003). Whereas Varimax rotation has the advantage of maintaining component orthogonality (Kayser & Tenke, 2006a), and obtaining simple structure has been proven useful in our prior work in temporal, spatial, spectral and time-frequency EEG domains, we recognize that oblique rotation procedures have yielded more favorable PCA outcomes in certain contexts (Barry & De Blasio, 2018; Dien, 2010; Scharf & Nestler, 2018). Although the effect of different blind source separation algorithms and rotation schemes on identification of RSNs is beyond the scope of this report, it will be an important question for future study.

Given the exploratory and developmental nature of the proposed CSD-fcPCA approach, we adopted a conservative “successive-hurdles” approach for selecting components. For step-one components, only components accounting for at least 1% of the total variance were included unless a factor needed to be considered for a priori reasons, which is in line with previous work (e.g., Kayser & Tenke, 2003; Tenke & Kayser, 2005). For these reasons, a midfrontal theta component that was below the 1% threshold was selected for our analysis because of *a priori* hypotheses concerning its potential as a candidate RSN (Cavanagh & Frank, 2014; Scheeringa et al., 2008; see Smith et al., 2020). For step-two components, only components that were evident across recording sessions, odd/even epochs, and in at least 2 out of the 4 acquisition sites were considered for further study.

To aid visualization and interpretation of network components, we opted to display only the top 10% of connectivity component loadings (249 loadings). These top component loadings were rank-ordered prior to visualization such that the strongest loadings had the largest ranks (e.g., 249 was the strongest link numerically). FC links can also be integrated across sensors to indicate node degree, a measure of the total number of functional connections between a particular sensor and every other sensor. In this case, the top 249 loadings were binarized, and then summed across sensors to produce a topography of node degree for each component. The resulting two types of FC component topography plots (i.e., edges and nodes strength) were evaluated in tandem to aid interpretation of identified RSNs. Importantly, our choice to display only the top 10% of loadings did not affect FC component quantification using the corresponding scores.

### 2.5 Statistics

Intraclass Correlations (ICCs) were used to evaluate internal consistency and test-retest reliability. Each participant was treated as a rater with two observations, operationalized as component scores from odd and even trials of session 1 (internal consistency), or all trials from session 1 and 2 (retest reliability), collapsed across EC/EO conditions. ICCs (1,k) were calculated from a one-way random analysis of variance (ANOVA; Shrout & Fleiss, 1979).

Tucker congruence coefficients (ϕ) were used to evaluate similarity in component loadings across PCA solutions computed on different subsamples of data (Lorenzo-Seva & ten Berge, 2006). These coefficients indicate to what extent two components are considered to be “equal” (ϕ ≥ .95) or to have “fair similarity” (.85 ≤ ϕ < .95).

Repeated measures ANOVA (SPSS GLM) was used to evaluate condition differences (EC/EO) separately for each RSN, with *p* < .05 considered statistically significant. However, effects of marginal significance (*p* < .10) are also reported. Effect sizes are given as Cohen’s *f* (Cohen, 1988).

## 3. Results

### 3.1.1 Spectral connectivity components (step one)

For the complete dataset, step one identified six spectral FC components with loading peaks at 4.2, 5.1, 6.3, 8.0, 10.2, and 13.0 Hz (Figure 1A). These components exhibited unambiguous peak loadings within canonical theta and alpha frequency bands (4-13 Hz) that were of primary interest and collectively explained 86.1% of spectral FC variance. Notably, the three alpha FC components peaking between 8 and 13 Hz explained the largest proportion of variance, which is consistent with the dwPLI grand mean revealing a distinct alpha peak with higher variability compared to the other frequencies (see gray line with shaded area in Figure 1A). These six components were used to spectrally filter FC data prior to the second PCA step.

Two additional step-one components with at least 1% explained variance had incomplete loadings with peaks at lower (3.3 Hz) and upper (16.0 Hz) bounds of the 3 to 16 Hz analysis window. Another component (9.0 Hz) was characterized by negative loadings suspicious for artifact from Varimax-PCA (i.e., adhering to the orthogonality requirement). Accordingly, these three components (Figure 1A) were not further considered.

All spectral FC PCA solutions (step one) separately obtained for each data subset (i.e., each session × epoch [odd/even] as well as for each data collection site) included 6 loadings that were highly comparable to those 6 obtained in step one for the combined data (Figure 1B; all ϕs ≥ .98), rendering these subset loadings effectively equal.

### 3.1.2 Spatial connectivity (network) components (step two)

Figures 2 and 3 show the step-two CSD-fcPCA solutions, displaying loading topographies (edges and nodes strength, respectively) for the first five spatial factors extracted (columns 2-6) for each of the 6 spectral FC components considered (rows). Step two identified ten alpha connectivity components (Figures 2 and 3, rows 1-3), each accounting for at least 1% of the total variance, and collectively accounted for 43.6%. However, two of the mid-alpha (10.2 Hz) and one of the low-alpha (8.0 Hz) spatial FC components meeting this criterion were found to be insufficiently robust across subgroup data sets, as detailed in the following section, and were therefore not further considered.

**Figure 2.**
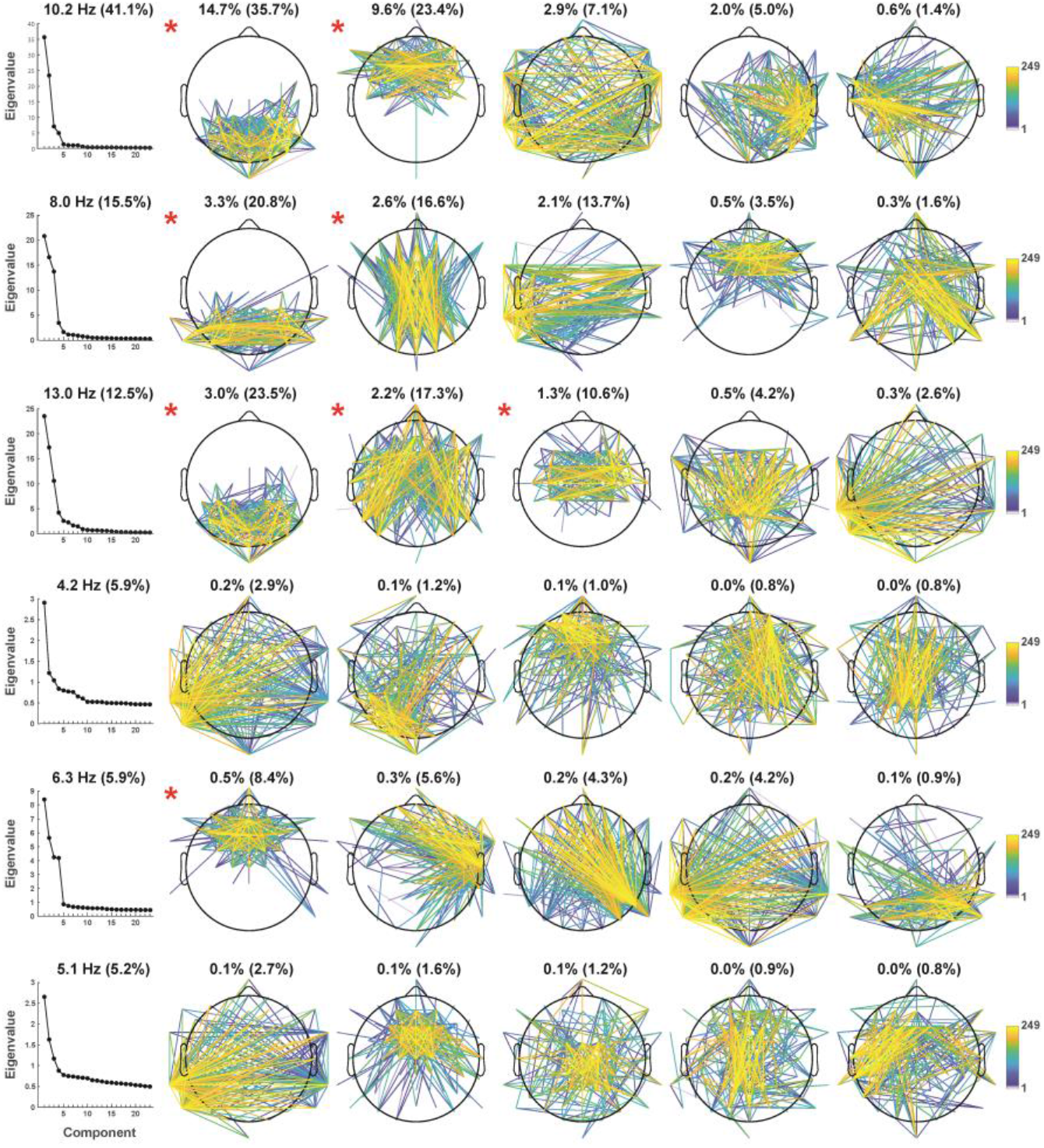
Edge strength of spatial FC components identified via step-two PCA. Connectivity strength between scalp sites was ranked and the top 10% of all edges are shown (249 corresponds to the highest rank). Column 1 depicts Eigenvalue plots (in percent of explained variance) of each step-two PCA solution, each subplot title indicates the corresponding step-one spectral component with its step-one explained variance (in parentheses). Columns 2-6 show the resulting loading topographies of the first 5 spatial components extracted, each subplot title lists the total variance explained and the corresponding step-two variance (in parentheses). Head maps marked with a red asterisk satisfied criteria for further analysis (see text).

**Figure 3.**
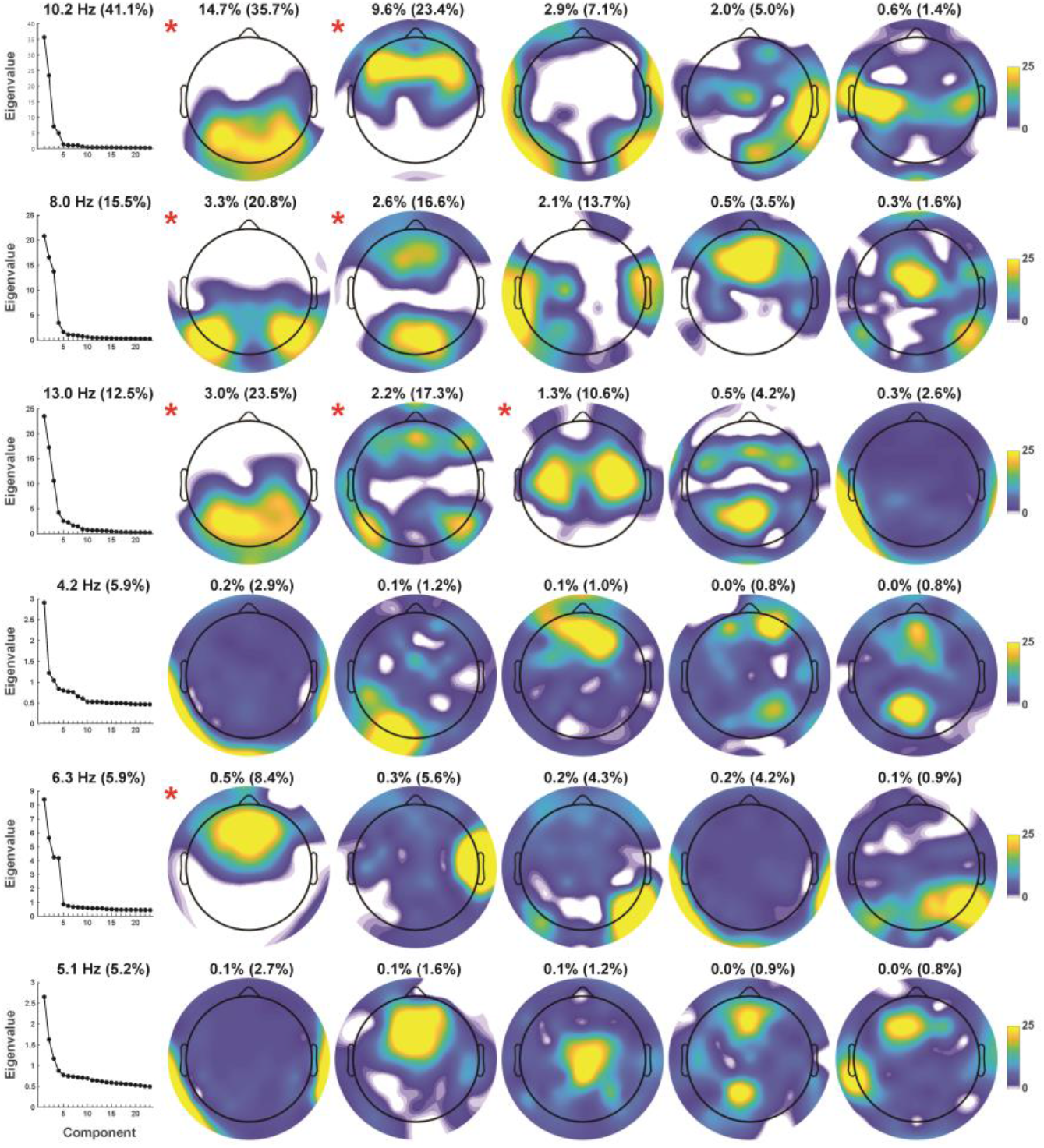
Node strength of spatial FC components identified via step-two PCA (organized as in Figure 2). Connectivity strength of each scalp site reflects the sum of the binarized top 10% loadings (as shown in Figure 2). The joint consideration of edge and node topographies allow for a concise interpretation of the resting state network (RSN) identified by each step-two FC component.

The retained spatial FC components are marked with a red asterisk next to their topo maps in Figures 2 and 3. At the mid-alpha frequency (10.2 Hz), two candidate RSNs were characterized by local connectivity between posterior sensors (posterior mid-alpha) and by dense FC between frontal sensors (frontal mid-alpha; Figures 2 and 3, row 1, columns 2-3). At the low-alpha frequency (8.0 Hz), two candidate RSNs were characterized by strong connectivity between lateral posterior sensors (posterior low-alpha) and between mid-occipital and mid-frontal sensors (mid posterior-anterior low-alpha; Figures 2 and 3, row 2, columns 2-3). At the high-alpha frequency (13.0 Hz), three candidate RSNs were characterized by connectivity between occipitoparietal sensors (posterior high-alpha), connectivity between lateral posterior and frontal sensors (lateral posterior-anterior high-alpha), and local connectivity bridging mid-central regions across both hemispheres (sensorimotor high-alpha; Figures 2 and 3, row 3, columns 2-4).

Compared to their alpha FC component counterparts, all step-two PCA solutions for the three step-one theta FC components had a more even variance spread between all extracted spatial FC components, as revealed by their Eigenvalue plots (Figure 2, rows 4-6, column 1), with no component explaining more than 1% of the total variance. However, a midfrontal component at 6.3 Hz accounted for an appreciable amount of step-two variance (8.4%), and was congruent with hypotheses regarding a midfrontal theta RSN (Figures 2 and 3, row 5, column 2). This midfrontal theta component was also identified in 7 of the 8 subgroup data sets (Tucker’s congruence coefficients, ϕ ≥ .86; see Supplemental Figure S5, column 1), suggesting FC component stability despite its low total variance. A peak frequency for midfrontal theta between 6 and 8 Hz, as opposed to a lower theta peak frequency, may also be more consistent with previous findings on midfrontal theta at rest (Kubicki, Herrmann, Fichte, & Freund, 1979; Pizzagalli, Oakes, & Davidson, 2003; Scheeringa et al., 2008).

Thus, a total of 8 RSN components (37.0% total FC variance explained) were retained for further analysis, seven of which had a frequency peak within the alpha band (36.5%) and one with a frequency peak in the theta band (0.5%).

### 3.2 Consistency of RSN component loadings

RSN loadings (step two) from aggregate data and subsamples demonstrated fair similarity and equality in most instances. For the eight RSNs considered for further analysis (i.e., red asterisk in Figures 2 and 3), all had corresponding RSNs (ϕ ≥ .89) in the 4 data subsets that included all participants (i.e., session [1/2] × epoch [odd/even]), and for at least two collection sites for which subsample sizes varied between *n* = 7 and *n* = 10 (see Supplementary Figures S1–S6 for edges and nodes strength topographies, explained variance, and Tucker’s congruence coefficients for all subsamples compared to the step-two PCA solutions of the entire dataset). The greatest correspondence between RSNs of the entire sample and the subsamples was seen for the first two RSNs of each alpha FC component and also the midfrontal theta FC component, all of which had 7 or 8 matches out of 8 subsamples (ϕ ≥ .88).

### 3.3 Reliability of RSN component scores

Split-half internal consistency for data collected at session one was excellent for all components (ICCs > .95). Test-retest ICCs showed moderate to excellent reliability across testing sessions for all components (.70 ≤ ICCs < .96; Koo & Li, 2016; Shrout & Fleiss, 1979). There was considerable similarity in RSN component scores for individual participants across sessions relative to the variability in scores between participants (Figure 4).

**Figure 4.**
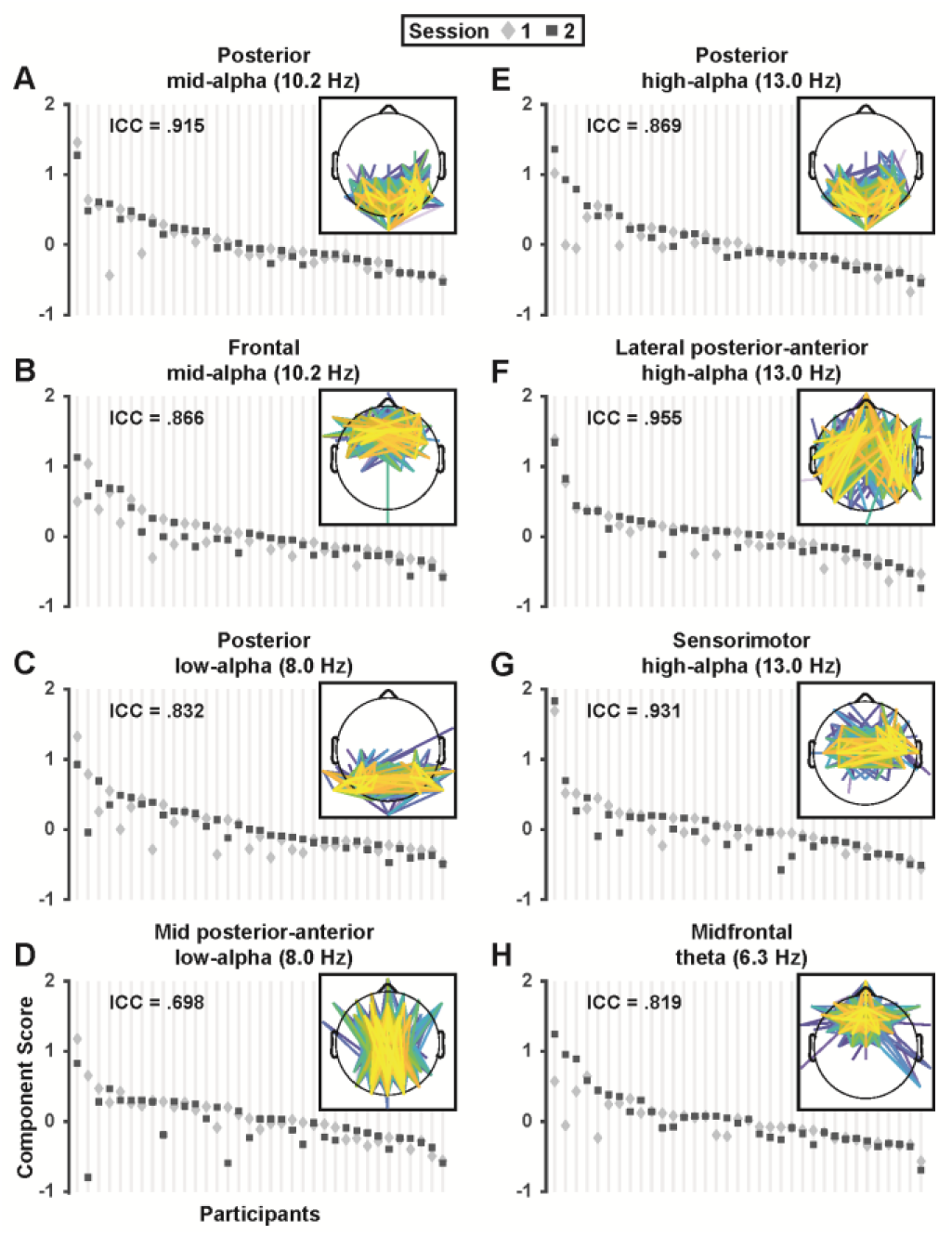
Test-retest reliability of mid-alpha (A, B), low-alpha (C, D), high-alpha (E-G) and theta (H) RSN component scores. For each RSN, mean component scores (pooled across EO/EC conditions and odd/even epochs) are depicted for each individual participant (*N* = 35; vertical lines) in session 1 (diamond) and 2 (square). For each component, participants were sorted by their maximum score across sessions to demonstrate the range of scores across participants relative to the individual variation in scores between session 1 and 2. The ICC value in upper left of each panel indicates the proportion of between-subjects versus between-sessions variance for individual components. Insets repeat RSN edge strength topographies depicted in Fig. 2.

### 3.4 Eyes open/closed condition differences

The posterior mid-alpha (8.0 Hz) RSN was greatly enhanced for EC compared to EO epochs, *F*(1,34) = 16.4, *p* = .0003, *f* = .695 (EC vs EO, Mean ±SD, .114 ±0.072 vs −.114 ±0.051; Figure 5A), and the posterior high-alpha (13.0 Hz) RSN was also greater for EC than EO, *F*(1,34) = 5.80, *p* = .02, *f* = .413 (.055 ±0.065 vs −.055 ±0.063; Figure 5B). Interestingly, the opposite effect (EO > EC) was seen for the two other 13.0 Hz RSNs, that is, the lateral posterior-anterior high-alpha RSN, *F*(1,34) = 3.74, *p* = .06, *f* = .332 (−.036 ±0.070 vs .036 ±0.064; Figure 5C), and the sensorimotor high-alpha RSN, *F*(1,34) = 4.71, *p* = .04, *f* = .372 (−.050 ±0.057 vs. 050 ±0.083; Figure 5D). There were no significant condition differences for any of the other RSNs (all *F*(1,34) ≤ 1.83, all *p* ≥ .18).

**Figure 5.**
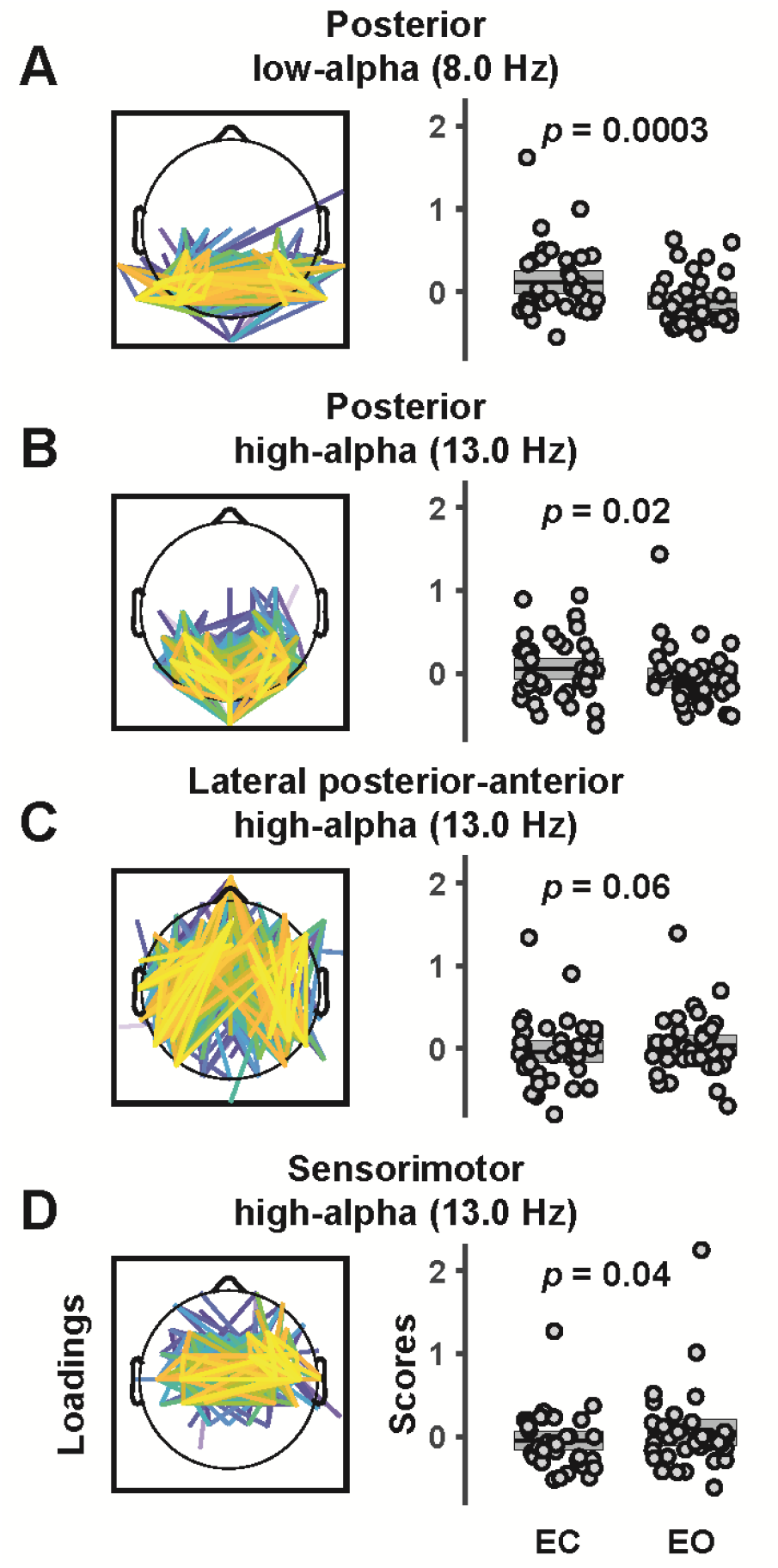
Low-alpha (A) and high-alpha (B-D) RSNs (spatial FC component loadings) showing a difference between eyes closed (EC) and eyes open (EO) conditions. Box plots show individual component scores for EC and EO (pooled across sessions and odd/even epochs) and sample means (N = 35) with the standard error of the mean (±SEM). Whereas posterior low-alpha and high-alpha FC components were stronger during EC than EO, lateral posterior-anterior and sensorimotor high-alpha RSNs were more pronounced for EO than EC conditions. Results demonstrate the functional specificity of network components between and within frequencies.

## 4. Discussion

The current findings present a comprehensive summary of phase-based RSNs intrinsic to resting EEG recordings using a reference-free data-driven approach. First, we minimized interpretational confounds due to volume conduction and choice of EEG recording reference (Bastos & Schoffelen, 2015; Tenke & Kayser, 2015) by using CSD transform as a spatial filter. Second, we employed a popular FC metric that is resistant to spurious connectivity estimates (dwPLI; Vinck et al., 2011) and has previously demonstrated adequate test-retest reliability for sensor-level EEG FC (Hardmeier et al., 2014). Third, a two-step PCA extending our prior work (e.g., Tenke & Kayser, 2005; Kayser & Tenke, 2006b) provided a concise and efficient summary of millions of data points in the form of eight distinct RSNs that explained over 37% of the connectivity variance in resting EEG data. Notably, only a targeted spectral subrange (3 – 16 Hz) was submitted to PCA, thus RSNs at lower (delta) or higher (beta, gamma) frequencies were not considered. However, there is no reason to suspect that the CSD-fcPCA approach developed here would not also work for other frequency ranges. These observations suggest that a generic linear approach to EEG analysis (Parra, Spence, Gerson, & Sajda, 2005) used to identify temporal (e.g., Tenke, Kayser, Stewart, & Bruder, 2010), spatial (e.g., Cohen, 2017; Dien, 2012; Tenke et al., 2008), spectral (e.g., Barry & De Blasio, 2018; Tenke & Kayser, 2005) and time-frequency (e.g., Kayser & Tenke, 2015a; Kayser et al., 2014) components can be extended towards the detection of reliable RSN components.

In most cases, RSNs were similar across data subsamples, suggesting that the identified RSNs were generalizable and not the result of a few isolated high-variance cases. Specifically, the identified RSNs were stable across recording sessions, internally consistent, and mostly comparable across collection sites. All RSNs demonstrated good-to-excellent internal consistency and one-week retest reliability. Whereas several RSNs converged with previous findings, there were a few unexpected alpha-band RSNs that had not been previously identified in resting EEG-dwPLI studies, but were nonetheless similar to FC patterns found in event-related designs. Altogether, we identified multiple candidate phase-based RSNs that were robust and reliable, strongly suggesting that CSD-fcPCA is a useful tool for the identification and quantification of RSNs in resting EEG recordings.

### 4.1 Findings and context

The present results align with previous work demonstrating the importance of alpha rhythms to RSNs (Chapeton, Haque, Wittig, Inati, & Zaghloul, 2019; Knyazev, Slobodskoj-Plusnin, Bocharov, & Pylkova, 2011; Sadaghiani & Kleinschmidt, 2016). Seven of the eight RSNs identified within a 3 – 16 Hz frequency range revealed a spectral peak within the canonical alpha band (8 – 13 Hz), with another RSN demonstrating a spectral peak at 6.3 Hz, well within the conventional theta band.

Three alpha components at differing peak frequencies (8.0, 10.2, and 13.0 Hz) were characterized by an abundance of connections between posterior sensors, consistent with typical alpha topographies (e.g., Smith et al., 2020). The fact that two out of three posterior alpha components demonstrated a significant condition difference underscores the existence of functionally dissociable alpha generators at distinct alpha frequency bands (e.g., Barry, De Blasio, Fogarty, & Clarke, 2020; Berger, 1929; Tenke & Kayser, 2005). A RSN with dense connections between sensors over sensorimotor cortex (sensorimotor high-alpha) was consistent with the well-known mu rhythm of the somatosensory system (e.g., Pfurtscheller et al., 1997)^6^. In addition, we identified a low alpha component (mid posterior-anterior low-alpha) having several long-range connections between mid-frontal and mid-occipital sensors that was similar to an 8 – 10 Hz pattern reported by Hardmeier et al. (2014), a 8-13 Hz midline pattern found by Hillebrand et al. (2016), and a midline FC component identified by Mehrkanoon et al. (2014).

We also found two additional and unexpected alpha RSNs: 1) a mid-alpha (10.2 Hz) component with local connections over frontal regions, and 2) a lateral posterior-anterior high-alpha (13.0 Hz) component characterized by long-range connections between lateral-frontal and lateral-parietal regions. Our impression is that CSD-fcPCA facilitated the detection of these novel RSNs that overlap in frequency and topography with other RSNs. For example, studies collapsing across one (Hillebrand et al., 2012; Stam et al., 2007) or two (Hardmeier et al., 2014; Hillebrand et al., 2016) alpha bands are by definition only capable of identifying RSNs within one or two alpha frequency bands, respectively. This finding suggests that multivariate approaches may be a necessary requisite for detecting RSNs with spectral-spatial overlap.

A subset of RSNs also varied in strength as a function of opening and closing of the eyes. Two alpha FC components (posterior low-alpha, posterior high-alpha) paralleled the classical Berger effect (EC > EO), whereas two other alpha components (lateral posterior-anterior high-alpha and sensorimotor high-alpha) showed the opposite condition effect (EO > EC). Greater RSN strength during EO is compatible with enhanced high-alpha FC across frontoparietal regions during working memory tasks (Sadaghiani & Kleinschmidt, 2016) and enhanced high-alpha FC across sensorimotor regions during motor initiation (Pfurtscheller et al., 1997). However, this is admittedly a speculative interpretation given that these effect sizes were notably weaker than those for EC > EO effects (i.e., medium rather than large), and identifying the precise functional significance of these novel RSN components will require further study.

A midfrontal theta (6.3 Hz) RSN identified here explained only a small fraction of connectivity variance in resting EEG, much less than explained by alpha network components, which is consistent with spectral amplitude findings for mid-frontal theta (e.g., Smith et al., 2020). Nonetheless, this FC component was internally consistent, test-retest reliable, and identified in 7 out of 8 data subsamples. This 6.3 Hz midfrontal theta component is also consistent with prior spectral analyses with regard to peak frequency and topography (e.g., Pizzagalli et al., 2003; Scheeringa et al., 2008). Altogether, the present findings showcase a diversity of reliable EEG-RSNs with varying peak frequencies, topographies, and putative functions that are ripe for additional investigation.

### 4.2 Limitations

Several limitations warrant consideration when interpreting these findings. First, the current study employed a conventional resting EEG paradigm with eyes open and closed recording periods as the only experimental manipulation. Although this is the de facto standard in the field, it severely limits interpretation of EEG connectivity, particularly when developing a new FC metric, as little is yet known how this variation affects rhythmic connections between regions, if at all. Because the present analysis was restricted to 3−16 Hz, RSNs at other frequency ranges were not considered; we encourage researchers to adopt the approach here to aid in the identification of RSNs at lower and higher frequencies. We have no reason to suspect this approach would not extend to other frequencies, but this remains an empirical question.

Second, EEG data were acquired in different EEG labs using disparate hard- and software necessitating a substantial effort to equate all critical recording aspects across acquisition sites before allowing a combined data analysis (Tenke et al., 2017). Given these constraints, it seems fairly remarkable that robust and interpretable RSNs emerged, suggesting that more homogeneous data sets may yield even more promising results.

Third, the CSD-fcPCA analysis reported here was based on a specific phase-lag FC metric, the debiased-weighted Phase-Lag Index (dwPLI; Vinck et al., 2011). However, many other FC metrics have been proposed and employed over the years, each with different properties and (dis-)advantages (e.g., Cohen, 2014). It is likely that different metrics will generate different FC components (e.g., Imperatoni et al., 2019), which will be a topic for future study. We would argue that the proposed CSD-fcPCA approach will also be effective in identifying and summarizing distinct FC variance stemming from other FC metrics. The hope is that this will also apply to causal (i.e., directional) FC metrics (e.g., Seth et al., 2015), however, this may require amendments to the specific CSD-fcPCA implementation developed here.

Fourth, the current FC data decomposition was applied at sensor as opposed to brain level. The advantages of a planar (two-dimensional) CSD transform (surface Laplacian) are that the obtained estimates of radial current flow at scalp are effectively model-independent (e.g., Tenke & Kayser, 2012) whereas all other three-dimensional generator source inverse solutions are dependent of their imposed biophysical constraints (e.g., Grave de Peralta Menendez et al., 2004; Pascual-Marqui et al., 1994). Still, EEG source localization techniques are useful if applied appropriately (e.g., Michel & He, 2019). Given that the quantification of EEG spectra at sensor and brain level via frequency PCA has demonstrated considerable similarities (Smith et al., 2020), it may be promising to apply the proposed fcPCA approach in concert with distributed inverses or other source localization techniques.

### 4.3 Conclusions

A comprehensive data-driven approach to analyze phase-based EEG functional connectivity — CSD-fcPCA — revealed candidate RSNs that were remarkably stable within and across participants. RSNs identified with CSD-fcPCA demonstrated good-to-excellent internal consistency and one-week test-retest reliability. The new CSD-fcPCA approach introduced here found multiple frequency-specific connectivity components that aligned with and extended previous work. Moreover, we found two additional candidate RSNs that would have been undetectable in previous reports that pooled FC metrics across canonical frequency bands. Alpha band RSNs dominated the resting EEG in the 3-16 Hz range, with 7 out of 8 network components identified between 8-13 Hz (i.e., classic alpha band). A theta (6.3 Hz) component demonstrated prominent connectivity amongst midfrontal sensors and was also reliable. Overall, these results suggest there are multiple phase-based RSNs in the resting EEG that demonstrate adequate reliability for consideration in future basic research and clinical applications.

## Acknowledgements

This research was funded by National Institute of Mental Health (NIMH) award MH115299 (JK). EEG data were obtained as part of the EMBARC study under NIMH awards MH092221 and MH092250.

## Author Contributions

**Ezra E. Smith**: Analysis; Software; Visualization; Writing, reviewing and editing. **Tarik S. Bel-Bahar**: Writing, reviewing and editing. **Jürgen Kayser**: Conceptualization; Data curation; Project administration; Resources; Supervision; Analysis; Software; Visualization; Writing, reviewing and editing.

## Supplementary Material

**Figure S1.**
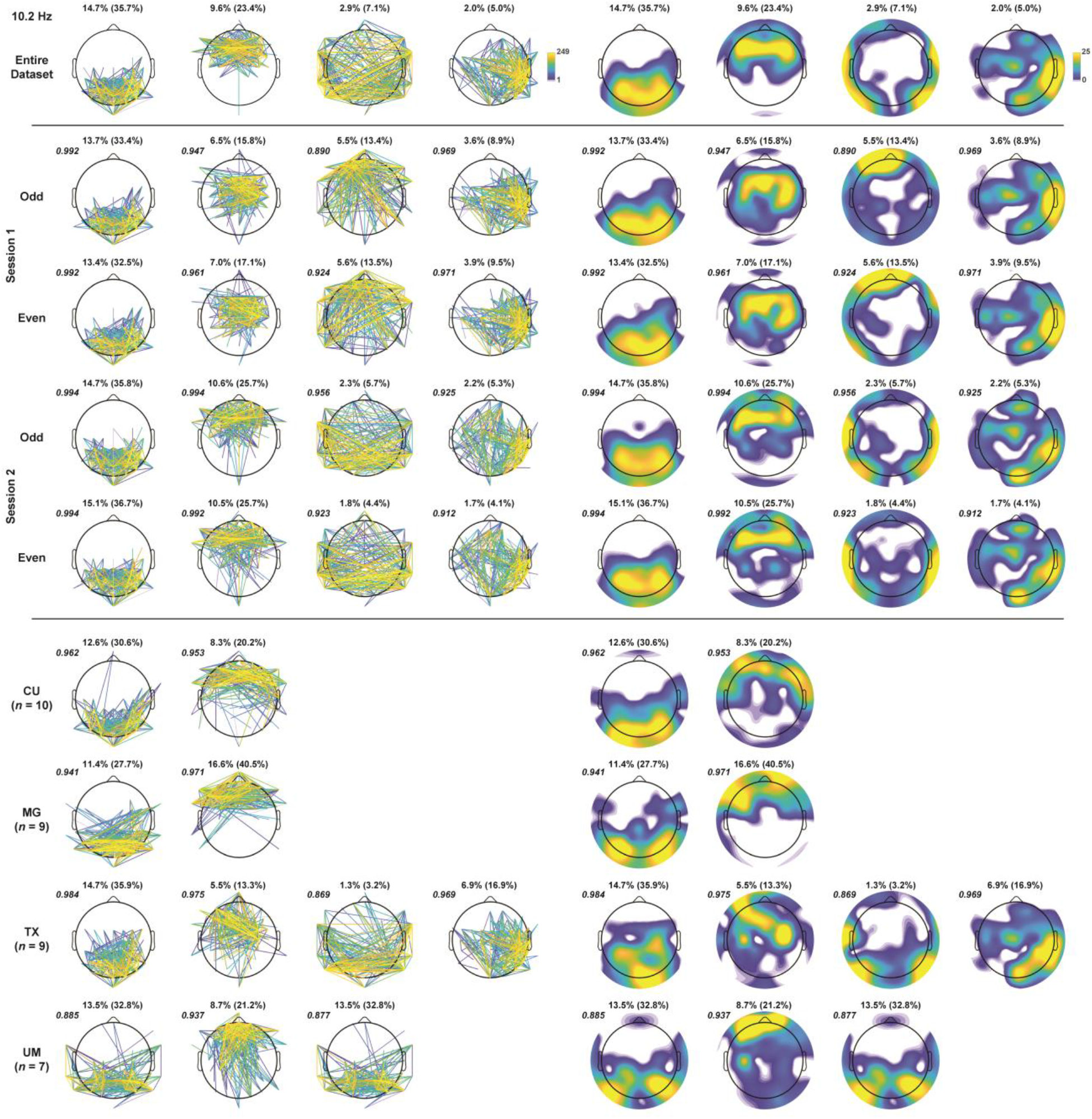
Edge and node strength of mid-alpha (10.2 Hz) spatial FC components identified in the step-two PCA applied to subsample data sets using step-one PCA components derived from the entire dataset (cf. Figures 2 and 3). For comparison, the top row shows edge and node strength topographies of the first 4 spatial FC components obtained for the original PCA solution. Rows depicts the best match for each subsample PCA solution, as determined by Tucker’s congruence coefficients ϕ (plotted in italics at the top left corner of each map); only loading maps having at least “fair similarity” (ϕ ≥ .85) are shown. Rows 2-5 show results for subsamples (*N* = 35) created by crossing session (1/2) with epoch (odd/even); rows 6-9 show those for subgroups based on EEG acquisition site (*n* = 7-10). Each subplot title lists the total variance explained and the corresponding step-two variance (in parentheses). Note that for subgroup UM (row 9), the same spatial FC component provided the best match for two different spatial FC components of the entire dataset (row 9, columns 1/5 and 3/7, respectively).

**Figure S2.**
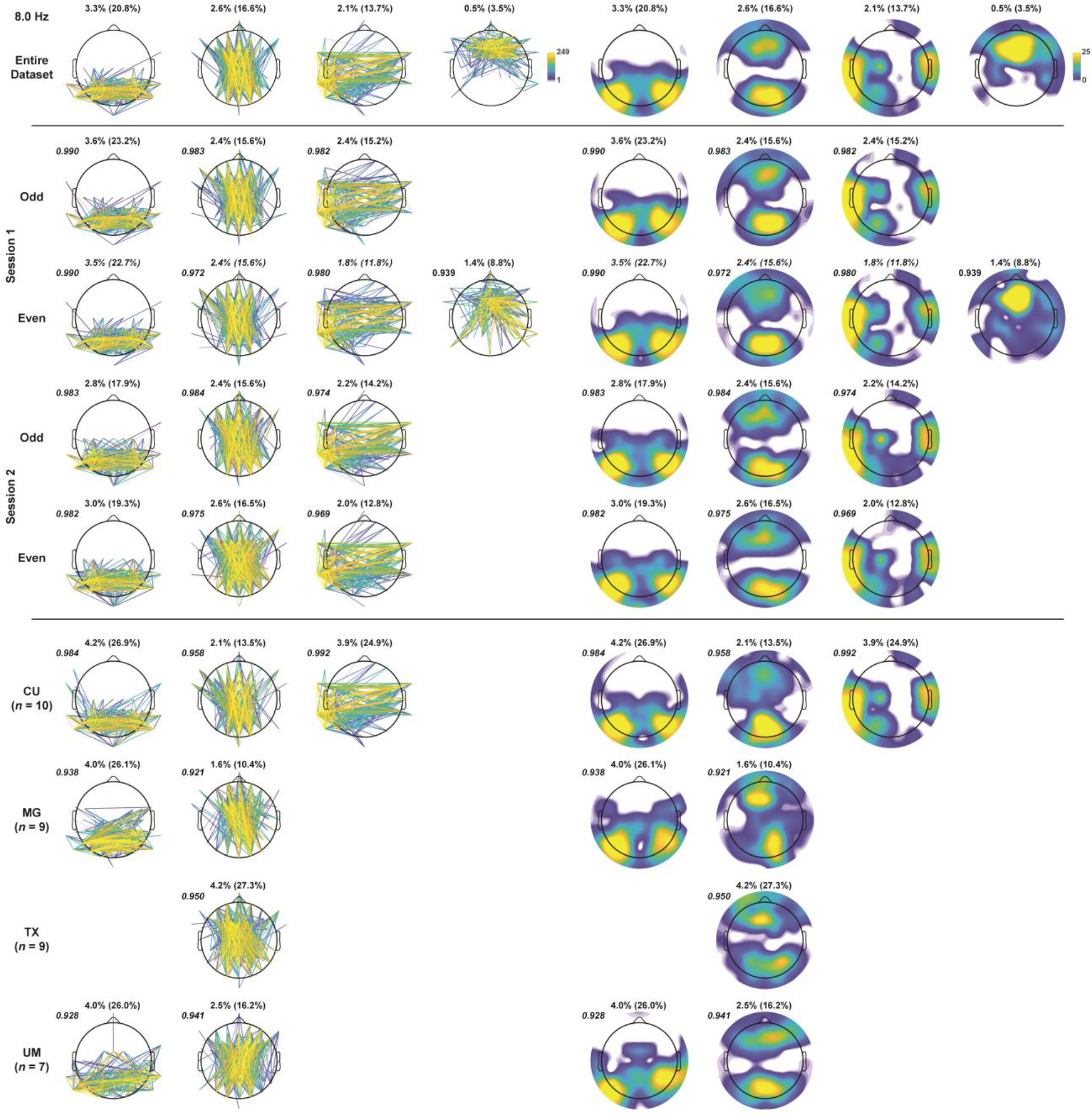
Edge and node strength of low-alpha (8.0 Hz) spatial FC components (all other details as in Figure S1).

**Figure S3.**
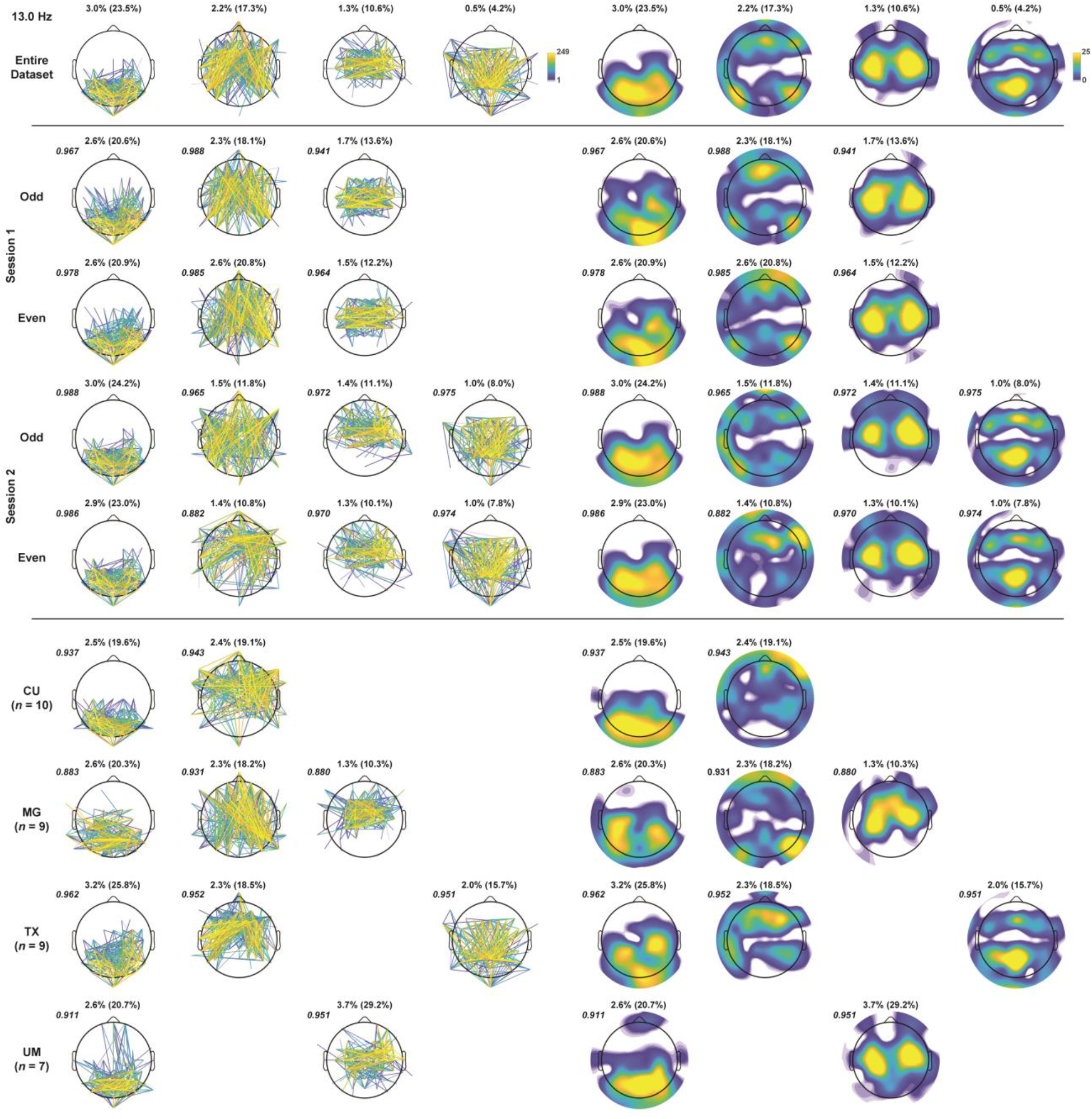
Edge and node strength of high-alpha (13.0 Hz) spatial FC components (all other details as in Figure S1).

**Figure S4.**
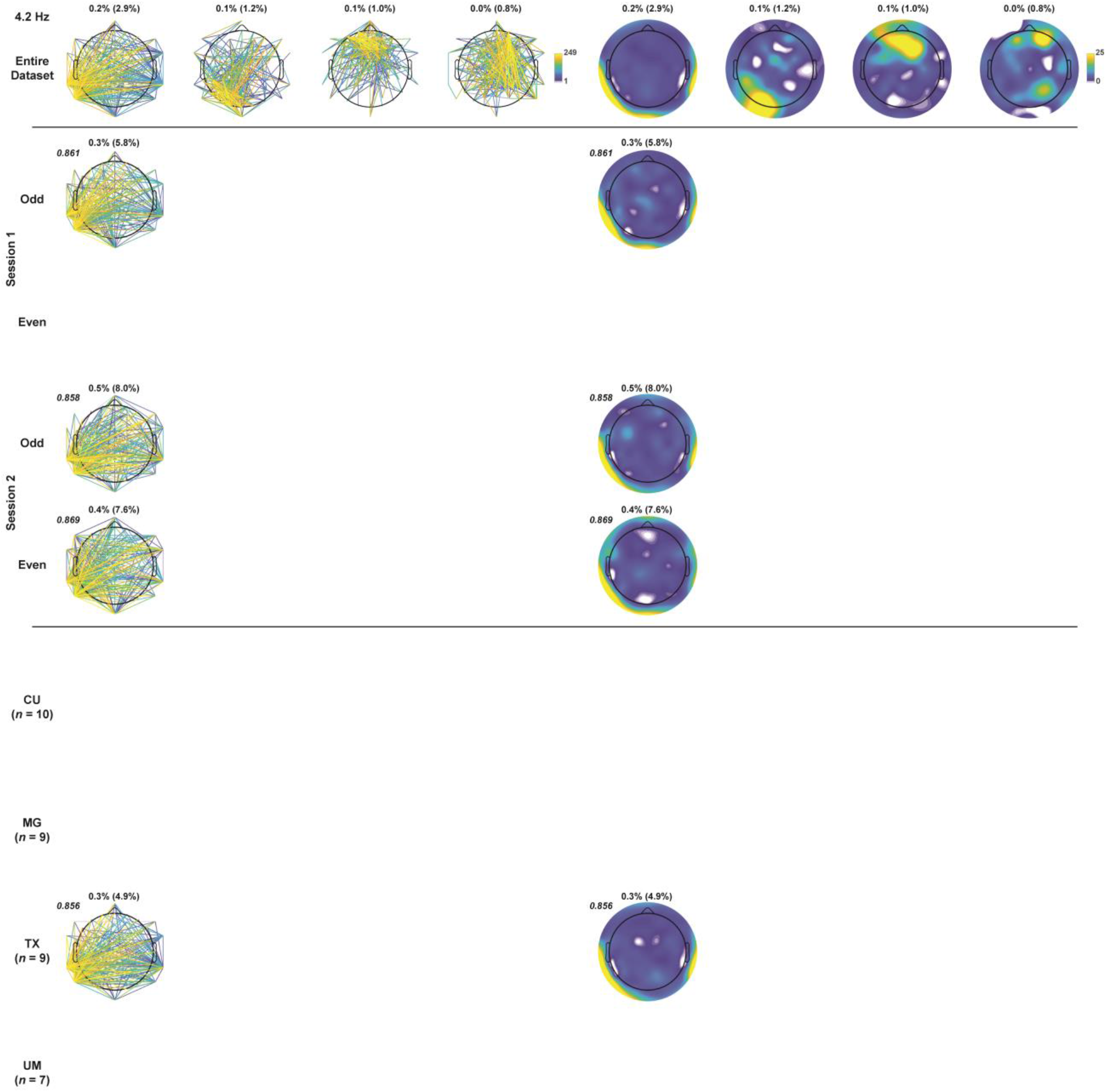
Edge and node strength of low-theta (4.2 Hz) spatial FC components (all other details as in Figure S1).

**Figure S5.**
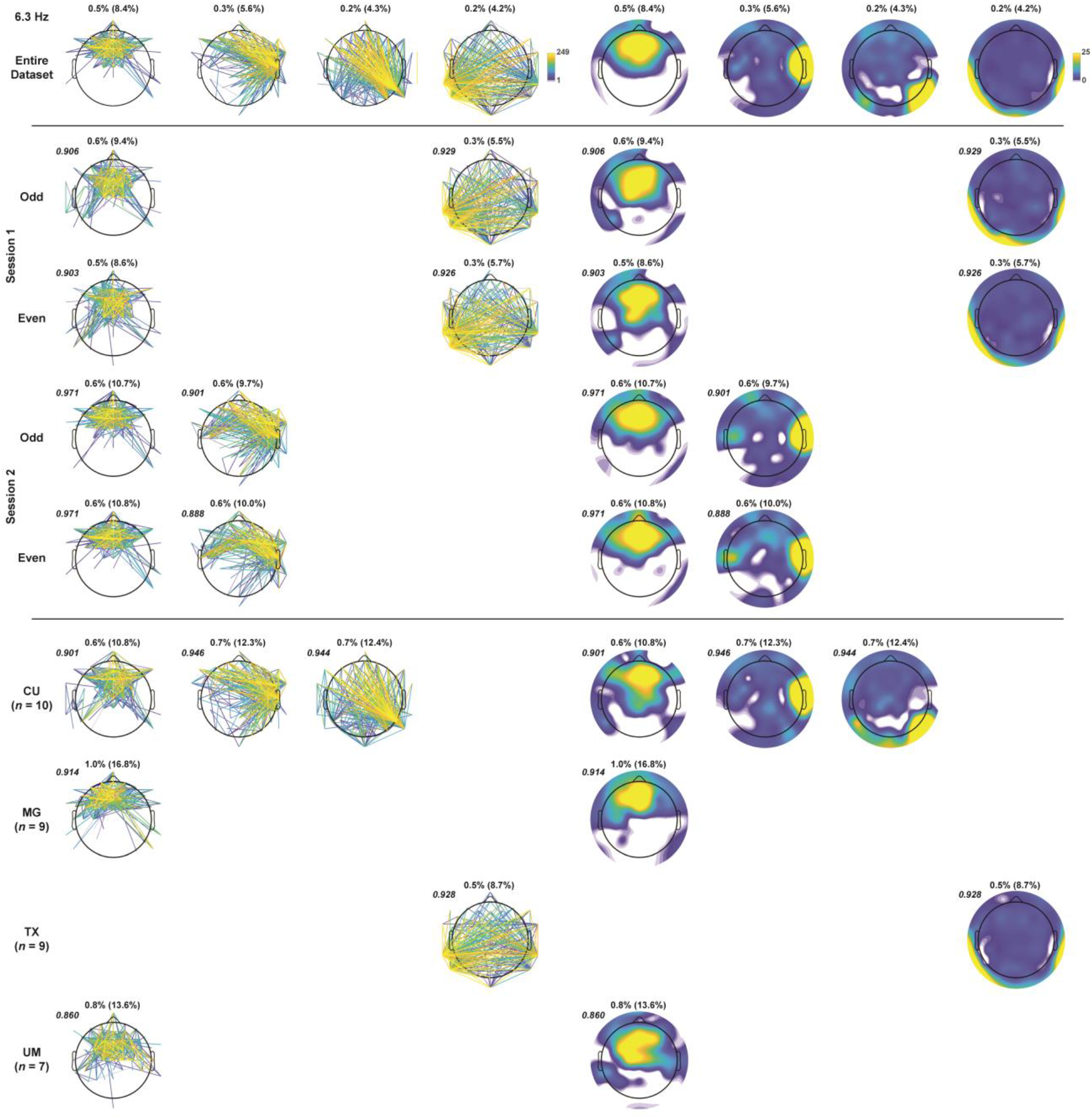
Edge and node strength of mid-theta (6.3 Hz) spatial FC components (all other details as in Figure S1).

**Figure S6.**
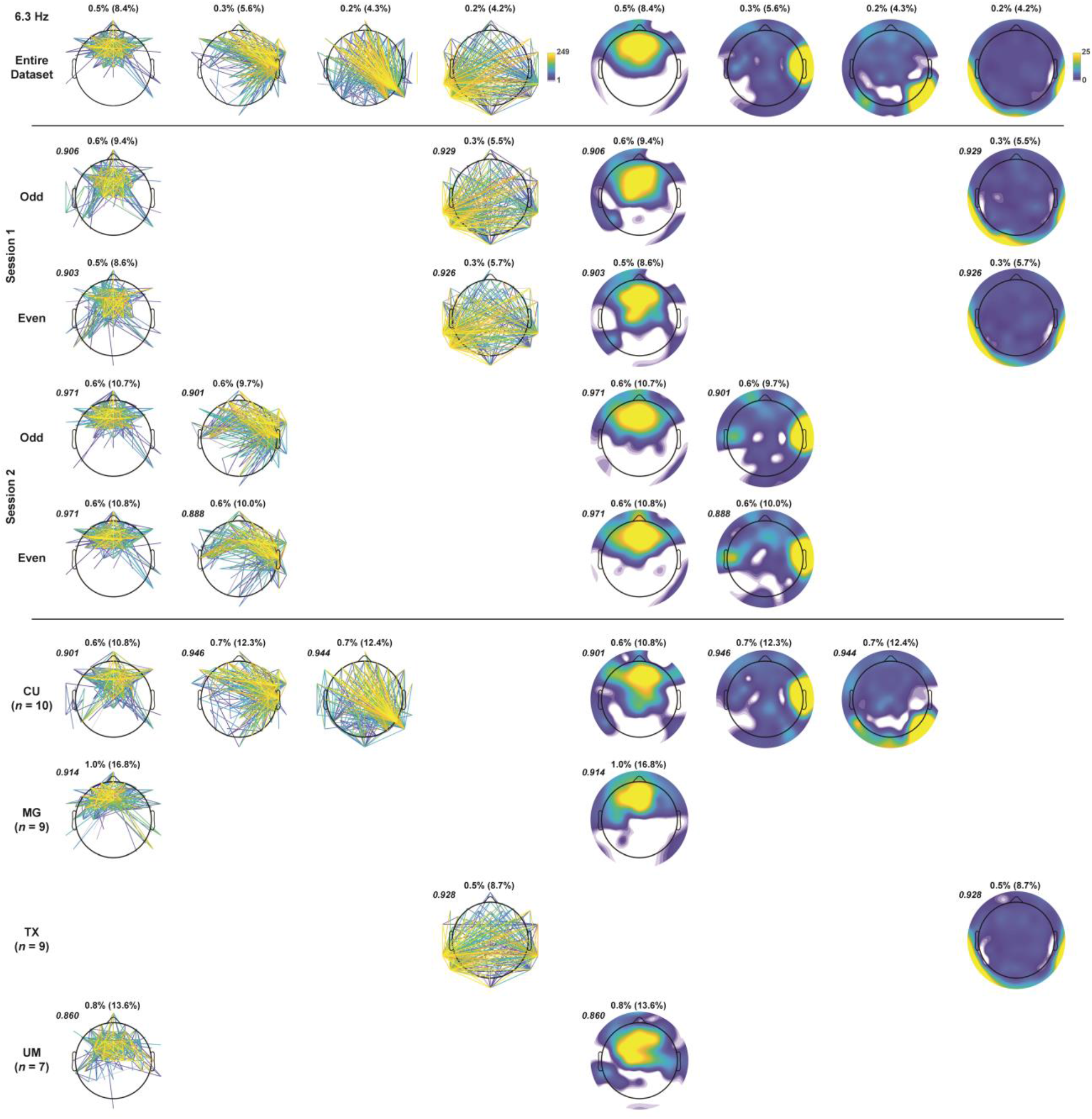
Edge and node strength of low/mid-theta (5.1 Hz) spatial FC components (all other details as in Figure S1).

## Footnotes

1 Compared to EEG, MEG studies have evidenced weaker test-retest reliability for global FC in both alpha and theta frequencies, particularly when using phase-based FC metrics (Candelaria-Cook, Schendel, Ojeda, Bustillo, & Stephen, 2020; Garces, Martin-Buro, & Maestu, 2016; Marquetand et al., 2019). It is not entirely clear why this is the case, but it may be partially attributed to differences in the types of dipoles (radial vs tangential) that are detected by EEG (both types) or MEG (only tangential), or to more FC resulting from volume-conduction and/or common sources in EEG recordings, in which case the “stronger” reliability would be misleading.

2 Informal pilot analyses suggested that adjacent CSD spline settings (e.g., *m* = 3 or 5) with default regularization (λ = 10^−5^) produced similar CSD-fcPCA results.

3 Morlet wavelets extracted 21 frequency bins (3, 3.3, 3.6, 3.9, 4.2, 4.6, 5, 5.4, 5.8, 6.4, 6.9, 7.5, 8.1, 8.8, 9.6, 10.4, 11.3, 12.3, 13.3, 14.5, 15.7 Hz) between 3 and 16 Hz which were interpolated to 42 logarithmically spaced bins (3, 3.1, 3.3, 3.4, 3.5, 3.7, 3.8, 4, 4.2, 4.3, 4.5, 4.7, 4.9, 5.1, 5.3, 5.5, 5.8, 6.0, 6.3, 6.5, 6.8, 7.1, 7.4, 7.7, 8.0, 8.3, 8.7, 9.0, 9.4, 9.8, 10.2, 10.6, 11.1, 11.5, 12, 12.5, 13, 13.6, 14.1, 14.7, 15.3, 16 Hz) to increase frequency resolution and the number of cases for the second PCA step.

4 Using the PCA of step one as a filter for the PCA of step two is notably different from other two-step PCA approaches that have submitted the associated factor scores of the step-one factors for step-two factorization (e.g., Dien, 2012). The approach here results in a step-two data matrix with a recommended cases-to-variables ratio (i.e., about 5:1; Gorsuch, 1983).

5 The grand mean, which is removed when using a covariance association matrix for PCA decomposition (e.g., Kayser & Tenke, 2003), was not reinserted during back-projection.

6 A low-alpha (8.0 Hz) bitemporal FC component that narrowly missed our stringent reliability criteria (see Supplementary Figure S2, columns 3 and 7) appeared to be consistent with the tau rhythm linked to supratemporal auditory cortex and sound suppression (e.g., Hari, Salmelin, Mäkelä, Salenius, & Helle, 1997; Tiihonen et al., 1991; Weisz et al., 2011; Yokosawa et al., 2020).

## References

Alagapan, S., Riddle, J., Huang, W.A., Hadar, E., Shin, H.W., & Frohlich, F. (2019). Network-Targeted, Multi-site Direct Cortical Stimulation Enhances Working Memory by Modulating Phase Lag of Low-Frequency Oscillations. Cell Rep, 29, 2590–2598 e2594. https://doi.org/10.1016/j.celrep.2019.10.072

Alschuler, D.M., Tenke, C.E., Bruder, G.E., & Kayser, J. (2014). Identifying electrode bridging from electrical distance distributions: a survey of publicly-available EEG data using a new method. Clinical Neurophysiology, 125, 484–490. https://doi.org/10.1016/j.clinph.2013.08.024

Barry, R.J., & De Blasio, F.M. (2018). EEG frequency PCA in EEG-ERP dynamics. Psychophysiology, 55, e13042. https://doi.org/10.1111/psyp.13042

Barry, R.J., De Blasio, F.M., Fogarty, J.S., & Clarke, A.R. (2020). Natural alpha frequency components in resting EEG and their relation to arousal. Clinical Neurophysiology, 131, 205–212. https://doi.org/10.1016/j.clinph.2019.10.018

Barry, R.J., De Blasio, F.M., & Karamacoska, D. (2019). Data-driven derivation of natural EEG frequency components: An optimised example assessing resting EEG in healthy ageing. Journal of Neuroscience Methods, 321, 1–11. https://doi.org/10.1016/j.jneumeth.2019.04.001

Bastos, A.M., & Schoffelen, J.M. (2015). A Tutorial Review of Functional Connectivity Analysis Methods and Their Interpretational Pitfalls. Front Syst Neurosci, 9, 175. https://doi.org/10.3389/fnsys.2015.00175

Berger, H. (1929). Über das Elektroenkephalogramm des Menschen. Archiv fur Psychiatrie und Nervenkrankheiten, 87, 527–570.

Biswal, B., Yetkin, F.Z., Haughton, V.M., & Hyde, J.S. (1995). Functional connectivity in the motor cortex of resting human brain using echo-planar MRI. Magnetic Resonance in Medicine, 34, 537–541. https://doi.org/10.1002/mrm.1910340409

Bressler, S.L., & Seth, A.K. (2011). Wiener-Granger causality: a well established methodology. Neuroimage, 58, 323–329. https://doi.org/10.1016/j.neuroimage.2010.02.059

Burle, B., Spieser, L., Roger, C., Casini, L., Hasbroucq, T., & Vidal, F. (2015). Spatial and temporal resolutions of EEG: Is it really black and white? A scalp current density view. International Journal of Psychophysiology, 97, 210–220. https://doi.org/10.1016/j.ijpsycho.2015.05.004

Candelaria-Cook, F.T., Schendel, M.E., Ojeda, C.J., Bustillo, J.R., & Stephen, J.M. (2020). Reduced parietal alpha power and psychotic symptoms: Test-retest reliability of resting-state magnetoencephalography in schizophrenia and healthy controls. Schizophrenia Research, 215, 229–240. https://doi.org/10.1016/j.schres.2019.10.023

Carvalhaes, C., & de Barros, J.A. (2015). The surface Laplacian technique in EEG: Theory and methods. International Journal of Psychophysiology, 97, 174–188. https://doi.org/10.1016/j.ijpsycho.2015.04.023

Cavanagh, J.F., & Frank, M.J. (2014). Frontal theta as a mechanism for cognitive control. Trends Cogn Sci, 18, 414–421. https://doi.org/10.1016/j.tics.2014.04.012

Chapeton, J.I., Haque, R., Wittig, J.H., Jr., Inati, S.K., & Zaghloul, K.A. (2019). Large-Scale Communication in the Human Brain Is Rhythmically Modulated through Alpha Coherence. Current Biology, 29, 2801–2811 e2805. https://doi.org/10.1016/j.cub.2019.07.014

Chapman, R.M., & McCrary, J.W. (1995). EP component identification and measurement by principal components analysis. Brain and Cognition, 27, 288–310. https://doi.org/10.1006/brcg.1995.1024

Cohen, J.D. (1988). Statistical power analysis for the behavioral sciences (2nd ed.). Hillsdale, NJ: Erlbaum.

Cohen, M.X. (2011). Error-related medial frontal theta activity predicts cingulate-related structural connectivity. Neuroimage, 55, 1373–1383. https://doi.org/10.1016/j.neuroimage.2010.12.072

Cohen, M.X. (2014). Analyzing neural time series data : theory and practice. Cambridge, MA: MIT Press.

Cohen, M.X. (2015). Comparison of different spatial transformations applied to EEG data: A case study of error processing. International Journal of Psychophysiology, 97, 245–257. https://doi.org/10.1016/j.ijpsycho.2014.09.013

Cohen, M.X. (2017). Comparison of linear spatial filters for identifying oscillatory activity in multichannel data. Journal of Neuroscience Methods, 278, 1–12. https://doi.org/10.1016/j.jneumeth.2016.12.016

Congedo, M., John, R.E., De Ridder, D., & Prichep, L. (2010). Group independent component analysis of resting state EEG in large normative samples. International Journal of Psychophysiology, 78, 89–99. https://doi.org/10.1016/j.ijpsycho.2010.06.003

Coquelet, N., De Tiege, X., Destoky, F., Roshchupkina, L., Bourguignon, M., Goldman, S., Peigneux, P., & Wens, V. (2020). Comparing MEG and high-density EEG for intrinsic functional connectivity mapping. Neuroimage, 210, 116556. https://doi.org/10.1016/j.neuroimage.2020.116556

Corlier, J., Wilson, A., Hunter, A.M., Vince-Cruz, N., Krantz, D., Levitt, J., Minzenberg, M.J., Ginder, N., Cook, I.A., & Leuchter, A.F. (2019). Changes in Functional Connectivity Predict Outcome of Repetitive Transcranial Magnetic Stimulation Treatment of Major Depressive Disorder. Cerebral Cortex, 29, 4958–4967. https://doi.org/10.1093/cercor/bhz035

David, O., Cosmelli, D., & Friston, K.J. (2004). Evaluation of different measures of functional connectivity using a neural mass model. Neuroimage, 21, 659–673. https://doi.org/10.1016/j.neuroimage.2003.10.006

Dien, J. (2010). Evaluating two-step PCA of ERP data with Geomin, Infomax, Oblimin, Promax, and Varimax rotations. Psychophysiology, 47, 170–183. https://doi.org/10.1111/j.1469-8986.2009.00885.x

Dien, J. (2012). Applying principal components analysis to event-related potentials: a tutorial. Dev Neuropsychol, 37, 497–517. https://doi.org/10.1080/87565641.2012.697503

Donchin, E. (1966). A multivariate approach to the analysis of average evoked potentials. IEEE Trans Biomed Eng, 13, 131–139. https://doi.org/10.1109/tbme.1966.4502423

Donchin, E., & Heffley, E.F. (1978). Multivariate analysis of evoked potential data: A tutorial review. In D.A. Otto (Ed.), Proceedings of the Fourth International Congress on Event-Related Slow Potentials of the Brain (EPIC IV), Hendersonville, NC, April 4-10, 1976 (pp. 555–572). Washington, DC: The Office.

Dunkin, J.J., Leuchter, A.F., Newton, T.F., & Cook, I.A. (1994). Reduced EEG coherence in dementia: state or trait marker? Biological Psychiatry, 35, 870–879. https://doi.org/10.1016/0006-3223(94)90023-x

Essl, M., & Rappelsberger, P. (1998). EEG coherence and reference signals: experimental results and mathematical explanations. Medical and Biological Engineering and Computing, 36, 399–406. https://doi.org/10.1007/BF02523206

Fein, G., Raz, J., Brown, F.F., & Merrin, E.L. (1988). Common reference coherence data are confounded by power and phase effects. Electroencephalography and Clinical Neurophysiology, 69, 581–584. https://doi.org/10.1016/0013-4694(88)90171-x

First, M.B., Spitzer, R.L., Gibbon, M., & Williams, J.B.W. (1996). Structured Clinical Interview for DSM-IV Axis-I Disorders – Non-patient Edition (SCID-NP). New York, NY: Biometrics Research Department, New York State Psychiatric Institute.

Forsyth, A., McMillan, R., Campbell, D., Malpas, G., Maxwell, E., Sleigh, J., Dukart, J., Hipp, J., & Muthukumaraswamy, S.D. (2020). Modulation of simultaneously collected hemodynamic and electrophysiological functional connectivity by ketamine and midazolam. Human Brain Mapping, 41, 1472–1494. https://doi.org/10.1002/hbm.24889

Fox, K.C.R., Foster, B.L., Kucyi, A., Daitch, A.L., & Parvizi, J. (2018). Intracranial Electrophysiology of the Human Default Network. Trends Cogn Sci, 22, 307–324. https://doi.org/10.1016/j.tics.2018.02.002

French, C.C., & Beaumont, J.G. (1984). A critical review of EEG coherence studies of hemisphere function. International Journal of Psychophysiology, 1, 241–254. https://doi.org/10.1016/0167-8760(84)90044-8

Fries, P. (2015). Rhythms for Cognition: Communication through Coherence. Neuron, 88, 220–235. https://doi.org/10.1016/j.neuron.2015.09.034

Gonzalez-Villar, A.J., Pidal-Miranda, M., Arias, M., Rodriguez-Salgado, D., & Carrillo-de-la-Pena, M.T. (2017). Electroencephalographic Evidence of Altered Top-Down Attentional Modulation in Fibromyalgia Patients During a Working Memory Task. Brain Topography, 30, 539–547. https://doi.org/10.1007/s10548-017-0561-3

Gorsuch, R.L. (2003). Factor analysis. Handbook of psychology: Research methods in psychology, Vol. 2. (pp. 143–164). Hoboken, NJ, US: John Wiley & Sons Inc.

Grave de Peralta Menendez, R., Murray, M.M., Michel, C.M., Martuzzi, R., & Gonzalez Andino, S.L. (2004). Electrical neuroimaging based on biophysical constraints. Neuroimage, 21, 527–539. https://doi.org/10.1016/j.neuroimage.2003.09.051

Hagemann, D., Naumann, E., & Thayer, J.F. (2001). The quest for the EEG reference revisited: a glance from brain asymmetry research. Psychophysiology, 38, 847–857.

Hardmeier, M., Hatz, F., Bousleiman, H., Schindler, C., Stam, C.J., & Fuhr, P. (2014). Reproducibility of functional connectivity and graph measures based on the phase lag index (PLI) and weighted phase lag index (wPLI) derived from high resolution EEG. PLoS ONE, 9, e108648. https://doi.org/10.1371/journal.pone.0108648

Hari, R., Salmelin, R., Makela, J.P., Salenius, S., & Helle, M. (1997). Magnetoencephalographic cortical rhythms. International Journal of Psychophysiology, 26, 51–62. https://doi.org/10.1016/s0167-8760(97)00755-1

Hillebrand, A., Barnes, G.R., Bosboom, J.L., Berendse, H.W., & Stam, C.J. (2012). Frequency-dependent functional connectivity within resting-state networks: an atlas-based MEG beamformer solution. Neuroimage, 59, 3909–3921. https://doi.org/10.1016/j.neuroimage.2011.11.005

Hillebrand, A., Tewarie, P., van Dellen, E., Yu, M., Carbo, E.W., Douw, L., Gouw, A.A., van Straaten, E.C., & Stam, C.J. (2016). Direction of information flow in large-scale resting-state networks is frequency-dependent. Proceedings of the National Academy of Sciences of the United States of America, 113, 3867–3872. https://doi.org/10.1073/pnas.1515657113

Hiltunen, T., Kantola, J., Abou Elseoud, A., Lepola, P., Suominen, K., Starck, T., Nikkinen, J., Remes, J., Tervonen, O., Palva, S., Kiviniemi, V., & Palva, J.M. (2014). Infra-slow EEG fluctuations are correlated with resting-state network dynamics in fMRI. Journal of Neuroscience, 34, 356–362. https://doi.org/10.1523/JNEUROSCI.0276-13.2014

Hipp, J.F., Hawellek, D.J., Corbetta, M., Siegel, M., & Engel, A.K. (2012). Large-scale cortical correlation structure of spontaneous oscillatory activity. Nature Neuroscience, 15, 884–890. https://doi.org/10.1038/nn.3101

Hjorth, B. (1975). An on-line transformation of EEG scalp potentials into orthogonal source derivations. Electroencephalography and Clinical Neurophysiology, 39, 526–530. https://doi.org/10.1016/0013-4694(75)90056-5

Holler, Y., Butz, K., Thomschewski, A., Schmid, E., Uhl, A., Bathke, A.C., Zimmermann, G., Tomasi, S.O., Nardone, R., Staffen, W., Holler, P., Leitinger, M., Hofler, J., Kalss, G., Taylor, A.C., Kuchukhidze, G., & Trinka, E. (2017). Reliability of EEG Interactions Differs between Measures and Is Specific for Neurological Diseases. Front Hum Neurosci, 11, 350. https://doi.org/10.3389/fnhum.2017.00350

Imperatori, L.S., Betta, M., Cecchetti, L., Canales-Johnson, A., Ricciardi, E., Siclari, F., Pietrini, P., Chennu, S., & Bernardi, G. (2019). EEG functional connectivity metrics wPLI and wSMI account for distinct types of brain functional interactions. Sci Rep, 9, 8894. https://doi.org/10.1038/s41598-019-45289-7

Jurcak, V., Tsuzuki, D., & Dan, I. (2007). 10/20, 10/10, and 10/5 systems revisited: their validity as relative head-surface-based positioning systems. Neuroimage, 34, 1600–1611. https://doi.org/10.1016/j.neuroimage.2006.09.024

Kayser, J. (2009). Current source density (CSD) interpolation using spherical splines – CSD Toolbox (Version 1.1). https://psychophysiology.cpmc.columbia.edu/software/CSDtoolbox/

Kayser, J., & Tenke, C.E. (2003). Optimizing PCA methodology for ERP component identification and measurement: theoretical rationale and empirical evaluation. Clinical Neurophysiology, 114, 2307–2325. https://doi.org/10.1016/s1388-2457(03)00241-4

Kayser, J., & Tenke, C.E. (2005). Trusting in or breaking with convention: towards a renaissance of principal components analysis in electrophysiology. Clinical Neurophysiology, 116, 1747–1753. https://doi.org/10.1016/j.clinph.2005.03.020

Kayser, J., & Tenke, C.E. (2006a). Consensus on PCA for ERP data, and sensibility of unrestricted solutions. Clinical Neurophysiology, 117, 703–707. https://doi.org/10.1016/j.clinph.2005.11.015

Kayser, J., & Tenke, C.E. (2006b). Principal components analysis of Laplacian waveforms as a generic method for identifying ERP generator patterns: I. Evaluation with auditory oddball tasks. Clinical Neurophysiology, 117, 348–368. https://doi.org/10.1016/j.clinph.2005.08.034

Kayser, J., & Tenke, C.E. (2006c). Principal components analysis of Laplacian waveforms as a generic method for identifying ERP generator patterns: II. Adequacy of low-density estimates. Clinical Neurophysiology, 117, 369–380. https://doi.org/10.1016/j.clinph.2005.08.033

Kayser, J., & Tenke, C.E. (2010). In search of the Rosetta Stone for scalp EEG: converging on reference-free techniques. Clinical Neurophysiology, 121, 1973–1975. https://doi.org/10.1016/j.clinph.2010.04.030

Kayser, J., & Tenke, C.E. (2015a). Hemifield-dependent N1 and event-related theta/delta oscillations: An unbiased comparison of surface Laplacian and common EEG reference choices. International Journal of Psychophysiology, 97, 258–270. https://doi.org/10.1016/j.ijpsycho.2014.12.011

Kayser, J., & Tenke, C.E. (2015b). Issues and considerations for using the scalp surface Laplacian in EEG/ERP research: A tutorial review. International Journal of Psychophysiology, 97, 189–209. https://doi.org/10.1016/j.ijpsycho.2015.04.012

Kayser, J., & Tenke, C.E. (2015c). On the benefits of using surface Laplacian (current source density) methodology in electrophysiology. International Journal of Psychophysiology, 97, 171–173. https://doi.org/10.1016/j.ijpsycho.2015.06.001

Kayser, J., Tenke, C.E., & Debener, S. (2000). Principal components analysis (PCA) as a tool for identifying EEG frequency bands: I. Methodological considerations and preliminary findings. Psychophysiology, 37, S54–S54.

Kayser, J., Tenke, C.E., Kroppmann, C.J., Alschuler, D.M., Fekri, S., Ben-David, S., Corcoran, C.M., & Bruder, G.E. (2014). Auditory event-related potentials and alpha oscillations in the psychosis prodrome: neuronal generator patterns during a novelty oddball task. International Journal of Psychophysiology, 91, 104–120. https://doi.org/10.1016/j.ijpsycho.2013.12.003

Knyazev, G.G., Slobodskoj-Plusnin, J.Y., Bocharov, A.V., & Pylkova, L.V. (2011). The default mode network and EEG alpha oscillations: an independent component analysis. Brain Research, 1402, 67–79. https://doi.org/10.1016/j.brainres.2011.05.052

Koo, T.K., & Li, M.Y. (2016). A Guideline of Selecting and Reporting Intraclass Correlation Coefficients for Reliability Research. J Chiropr Med, 15, 155–163. https://doi.org/10.1016/j.jcm.2016.02.012

Kubicki, S., Herrmann, W.M., Fichte, K., & Freund, G. (1979). Reflections on the topics: EEG frequency bands and regulation of vigilance. Pharmakopsychiatrie Neuro-Psychopharmakologie, 12, 237–245. https://doi.org/10.1055/s-0028-1094615

Kucyi, A., Schrouff, J., Bickel, S., Foster, B.L., Shine, J.M., & Parvizi, J. (2018). Intracranial Electrophysiology Reveals Reproducible Intrinsic Functional Connectivity within Human Brain Networks. Journal of Neuroscience, 38, 4230–4242. https://doi.org/10.1523/JNEUROSCI.0217-18.2018

Kuntzelman, K., & Miskovic, V. (2017). Reliability of graph metrics derived from resting-state human EEG. Psychophysiology, 54, 51–61. https://doi.org/10.1111/psyp.12600

Lachaux, J.P., Rodriguez, E., Martinerie, J., & Varela, F.J. (1999). Measuring phase synchrony in brain signals. Human Brain Mapping, 8, 194–208. https://doi.org/10.1002/(sici)1097-0193(1999)8:4<194::aid-hbm4>3.0.co;2-c

Lee, M.H., Smyser, C.D., & Shimony, J.S. (2013). Resting-state fMRI: a review of methods and clinical applications. AJNR: American Journal of Neuroradiology, 34, 1866–1872. https://doi.org/10.3174/ajnr.A3263

Leocani, L., & Comi, G. (1999). EEG coherence in pathological conditions. Journal of Clinical Neurophysiology, 16, 548–555. https://doi.org/10.1097/00004691-199911000-00006

Lorenzo-Seva, U., & ten Berge, J.M.F. (2006). Tucker’s Congruence Coefficient as a Meaningful Index of Factor Similarity. Methodology, 2, 57–64. https://doi.org/10.1027/1614-2241.2.2.57

Mahjoory, K., Nikulin, V.V., Botrel, L., Linkenkaer-Hansen, K., Fato, M.M., & Haufe, S. (2017). Consistency of EEG source localization and connectivity estimates. Neuroimage, 152, 590–601. https://doi.org/10.1016/j.neuroimage.2017.02.076

Mansour, L.S., Tian, Y., Yeo, B.T.T., Cropley, V., & Zalesky, A. (2021). High-resolution connectomic fingerprints: Mapping neural identity and behavior. Neuroimage, 229, 117695. https://doi.org/10.1016/j.neuroimage.2020.117695

Mantini, D., Perrucci, M.G., Del Gratta, C., Romani, G.L., & Corbetta, M. (2007). Electrophysiological signatures of resting state networks in the human brain. Proceedings of the National Academy of Sciences of the United States of America, 104, 13170–13175. https://doi.org/10.1073/pnas.0700668104

Marquetand, J., Vannoni, S., Carboni, M., Li Hegner, Y., Stier, C., Braun, C., & Focke, N.K. (2019). Reliability of Magnetoencephalography and High-Density Electroencephalography Resting-State Functional Connectivity Metrics. Brain Connect, 9, 539–553. https://doi.org/10.1089/brain.2019.0662

Martin-Buro, M.C., Garces, P., & Maestu, F. (2016). Test-retest reliability of resting-state magnetoencephalography power in sensor and source space. Human Brain Mapping, 37, 179–190. https://doi.org/10.1002/hbm.23027

Marzetti, L., Basti, A., Chella, F., D’Andrea, A., Syrjala, J., & Pizzella, V. (2019). Brain Functional Connectivity Through Phase Coupling of Neuronal Oscillations: A Perspective From Magnetoencephalography. Front Neurosci, 13, 964. https://doi.org/10.3389/fnins.2019.00964

Mehrkanoon, S., Breakspear, M., & Boonstra, T.W. (2014). Low-dimensional dynamics of resting-state cortical activity. Brain Topography, 27, 338–352. https://doi.org/10.1007/s10548-013-0319-5

Michel, C.M., & He, B. (2019). EEG source localization. Handb Clin Neurol, 160, 85–101. https://doi.org/10.1016/B978-0-444-64032-1.00006-0

NeuroScan Inc. (2003). SCAN 4.3 – Vol. II. EDIT 4.3 – Offline analysis of acquired data (Document number 2203, Revision D).

Nolte, G., Bai, O., Wheaton, L., Mari, Z., Vorbach, S., & Hallett, M. (2004). Identifying true brain interaction from EEG data using the imaginary part of coherency. Clinical Neurophysiology, 115, 2292–2307. https://doi.org/10.1016/j.clinph.2004.04.029

Nunez, P.L., Srinivasan, R., Westdorp, A.F., Wijesinghe, R.S., Tucker, D.M., Silberstein, R.B., & Cadusch, P.J. (1997). EEG coherency. I: Statistics, reference electrode, volume conduction, Laplacians, cortical imaging, and interpretation at multiple scales. Electroencephalography and Clinical Neurophysiology, 103, 499–515. https://doi.org/10.1016/s0013-4694(97)00066-7

Oldfield, R.C. (1971). The assessment and analysis of handedness: the Edinburgh inventory. Neuropsychologia, 9, 97–113. https://doi.org/10.1016/0028-3932(71)90067-4

Pan, W.J., Thompson, G.J., Magnuson, M.E., Jaeger, D., & Keilholz, S. (2013). Infraslow LFP correlates to resting-state fMRI BOLD signals. Neuroimage, 74, 288–297. https://doi.org/10.1016/j.neuroimage.2013.02.035

Parra, L.C., Spence, C.D., Gerson, A.D., & Sajda, P. (2005). Recipes for the linear analysis of EEG. Neuroimage, 28, 326–341. https://doi.org/10.1016/j.neuroimage.2005.05.032

Pascual-Marqui, R.D., Michel, C.M., & Lehmann, D. (1994). Low resolution electromagnetic tomography: a new method for localizing electrical activity in the brain. International Journal of Psychophysiology, 18, 49–65. https://doi.org/10.1016/0167-8760(84)90014-x

Perrin, F., Pernier, J., Bertrand, O., & Echallier, J.F. (1989). Spherical splines for scalp potential and current density mapping. Electroencephalography and Clinical Neurophysiology, 72, 184–187. [Corrigenda EEG 02274, EEG Clin. Neurophysiol., 1990, 76, 565] https://doi.org/10.1016/0013-4694(89)90180-6

Pfurtscheller, G., Neuper, C., Andrew, C., & Edlinger, G. (1997). Foot and hand area mu rhythms. International Journal of Psychophysiology, 26, 121–135. https://doi.org/10.1016/s0167-8760(97)00760-5

Pizzagalli, D.A. (2011). Frontocingulate dysfunction in depression: toward biomarkers of treatment response. Neuropsychopharmacology, 36, 183–206. https://doi.org/10.1038/npp.2010.166

Pizzagalli, D.A., Oakes, T.R., & Davidson, R.J. (2003). Coupling of theta activity and glucose metabolism in the human rostral anterior cingulate cortex: an EEG/PET study of normal and depressed subjects. Psychophysiology, 40, 939–949. https://doi.org/10.1111/1469-8986.00112

Portoles, O., Borst, J.P., & van Vugt, M.K. (2018). Characterizing synchrony patterns across cognitive task stages of associative recognition memory. European Journal of Neuroscience, 48, 2759–2769. https://doi.org/10.1111/ejn.13817

Raichle, M.E. (2015). The brain’s default mode network. Annual Review of Neuroscience, 38, 433–447. https://doi.org/10.1146/annurev-neuro-071013-014030

Raichle, M.E., MacLeod, A.M., Snyder, A.Z., Powers, W.J., Gusnard, D.A., & Shulman, G.L. (2001). A default mode of brain function. Proceedings of the National Academy of Sciences of the United States of America, 98, 676–682. https://doi.org/10.1073/pnas.98.2.676

Rizkallah, J., Amoud, H., Fraschini, M., Wendling, F., & Hassan, M. (2020). Exploring the Correlation Between M/EEG Source-Space and fMRI Networks at Rest. Brain Topography, 33, 151–160. https://doi.org/10.1007/s10548-020-00753-w

Roberts, A., Fillmore, P., & Decker, S. (2016). Clinical Applicability of the Test-retest Reliability of qEEG Coherence. NeuroRegulation, 3, 7–22. https://doi.org/10.15540/nr.3.1.7

Rouhinen, S., Siebenhuhner, F., Palva, J.M., & Palva, S. (2020). Spectral and Anatomical Patterns of Large-Scale Synchronization Predict Human Attentional Capacity. Cerebral Cortex, 30, 5293–5308. https://doi.org/10.1093/cercor/bhaa110

Sadaghiani, S., & Kleinschmidt, A. (2016). Brain Networks and alpha-Oscillations: Structural and Functional Foundations of Cognitive Control. Trends Cogn Sci, 20, 805–817. https://doi.org/10.1016/j.tics.2016.09.004

Sadaghiani, S., Scheeringa, R., Lehongre, K., Morillon, B., Giraud, A.L., & Kleinschmidt, A. (2010). Intrinsic connectivity networks, alpha oscillations, and tonic alertness: a simultaneous electroencephalography/functional magnetic resonance imaging study. Journal of Neuroscience, 30, 10243–10250. https://doi.org/10.1523/JNEUROSCI.1004-10.2010

Scharf, F., & Nestler, S. (2018). Principles behind variance misallocation in temporal exploratory factor analysis for ERP data: Insights from an inter-factor covariance decomposition. International Journal of Psychophysiology, 128, 119–136. https://doi.org/10.1016/j.ijpsycho.2018.03.019

Scheeringa, R., Bastiaansen, M.C., Petersson, K.M., Oostenveld, R., Norris, D.G., & Hagoort, P. (2008). Frontal theta EEG activity correlates negatively with the default mode network in resting state. International Journal of Psychophysiology, 67, 242–251. https://doi.org/10.1016/j.ijpsycho.2007.05.017

Seth, A.K., Barrett, A.B., & Barnett, L. (2015). Granger causality analysis in neuroscience and neuroimaging. Journal of Neuroscience, 35, 3293–3297. https://doi.org/10.1523/JNEUROSCI.4399-14.2015

Shrout, P.E., & Fleiss, J.L. (1979). Intraclass correlations: uses in assessing rater reliability. Psychological Bulletin, 86, 420–428. https://doi.org/10.1037//0033-2909.86.2.420

Siems, M., & Siegel, M. (2020). Dissociated neuronal phase- and amplitude-coupling patterns in the human brain. Neuroimage, 209, 116538. https://doi.org/10.1016/j.neuroimage.2020.116538

Smith, E.E., Reznik, S.J., Stewart, J.L., & Allen, J.J. (2017). Assessing and conceptualizing frontal EEG asymmetry: An updated primer on recording, processing, analyzing, and interpreting frontal alpha asymmetry. International Journal of Psychophysiology, 111, 98–114. https://doi.org/10.1016/j.ijpsycho.2016.11.005

Smith, E.E., Tenke, C.E., Deldin, P.J., Trivedi, M.H., Weissman, M.M., Auerbach, R.P., Bruder, G.E., Pizzagalli, D.A., & Kayser, J. (2020). Frontal theta and posterior alpha in resting EEG: A critical examination of convergent and discriminant validity. Psychophysiology, 57, e13483. https://doi.org/10.1111/psyp.13483

Stam, C.J., Nolte, G., & Daffertshofer, A. (2007). Phase lag index: assessment of functional connectivity from multi channel EEG and MEG with diminished bias from common sources. Human Brain Mapping, 28, 1178–1193. https://doi.org/10.1002/hbm.20346

Tenke, C.E., & Kayser, J. (2005). Reference-free quantification of EEG spectra: combining current source density (CSD) and frequency principal components analysis (fPCA). Clinical Neurophysiology, 116, 2826–2846. https://doi.org/10.1016/j.clinph.2005.08.007

Tenke, C.E., & Kayser, J. (2012). Generator localization by current source density (CSD): implications of volume conduction and field closure at intracranial and scalp resolutions. Clinical Neurophysiology, 123, 2328–2345. https://doi.org/10.1016/j.clinph.2012.06.005

Tenke, C.E., & Kayser, J. (2015). Surface Laplacians (SL) and phase properties of EEG rhythms: Simulated generators in a volume-conduction model. International Journal of Psychophysiology, 97, 285–298. https://doi.org/10.1016/j.ijpsycho.2015.05.008

Tenke, C.E., Kayser, J., Manna, C.G., Fekri, S., Kroppmann, C.J., Schaller, J.D., Alschuler, D.M., Stewart, J.W., McGrath, P.J., & Bruder, G.E. (2011). Current source density measures of electroencephalographic alpha predict antidepressant treatment response. Biological Psychiatry, 70, 388–394. https://doi.org/10.1016/j.biopsych.2011.02.016

Tenke, C.E., Kayser, J., Pechtel, P., Webb, C.A., Dillon, D.G., Goer, F., Murray, L., Deldin, P., Kurian, B.T., McGrath, P.J., Parsey, R., Trivedi, M., Fava, M., Weissman, M.M., McInnis, M., Abraham, K., J. E.A., Alschuler, D.M., Cooper, C., Pizzagalli, D.A., & Bruder, G.E. (2017). Demonstrating test-retest reliability of electrophysiological measures for healthy adults in a multisite study of biomarkers of antidepressant treatment response. Psychophysiology, 54, 34–50. https://doi.org/10.1111/psyp.12758

Tenke, C.E., Kayser, J., Shankman, S.A., Griggs, C.B., Leite, P., Stewart, J.W., & Bruder, G.E. (2008). Hemispatial PCA dissociates temporal from parietal ERP generator patterns: CSD components in healthy adults and depressed patients during a dichotic oddball task. International Journal of Psychophysiology, 67, 1–16. https://doi.org/10.1016/j.ijpsycho.2007.09.001

Tenke, C.E., Kayser, J., Stewart, J.W., & Bruder, G.E. (2010). Novelty P3 reductions in depression: characterization using principal components analysis (PCA) of current source density (CSD) waveforms. Psychophysiology, 47, 133–146. https://doi.org/10.1111/j.1469-8986.2009.00880.x

Tiihonen, J., Hari, R., Kajola, M., Karhu, J., Ahlfors, S., & Tissari, S. (1991). Magnetoencephalographic 10-Hz rhythm from the human auditory cortex. Neuroscience Letters, 129, 303–305. https://doi.org/10.1016/0304-3940(91)90486-d

Trivedi, M.H., McGrath, P.J., Fava, M., Parsey, R.V., Kurian, B.T., Phillips, M.L., Oquendo, M.A., Bruder, G., Pizzagalli, D., Toups, M., Cooper, C., Adams, P., Weyandt, S., Morris, D.W., Grannemann, B.D., Ogden, R.T., Buckner, R., McInnis, M., Kraemer, H.C., Petkova, E., Carmody, T.J., & Weissman, M.M. (2016). Establishing moderators and biosignatures of antidepressant response in clinical care (EMBARC): Rationale and design. Journal of Psychiatric Research, 78, 11–23. https://doi.org/10.1016/j.jpsychires.2016.03.001

Uhlhaas, P.J., Liddle, P., Linden, D.E.J., Nobre, A.C., Singh, K.D., & Gross, J. (2017). Magnetoencephalography as a Tool in Psychiatric Research: Current Status and Perspective. Biol Psychiatry Cogn Neurosci Neuroimaging, 2, 235–244. https://doi.org/10.1016/j.bpsc.2017.01.005

Van Boxtel, G.J.M. (1998). Computational and statistical methods for analyzing event-related potential data. Behavior Research Methods, Instruments, & Computers, 30, 87–102. https://doi.org/10.3758/bf03209419

van der Velde, B., Haartsen, R., & Kemner, C. (2019). Test-retest reliability of EEG network characteristics in infants. Brain Behav, 9, e01269. https://doi.org/10.1002/brb3.1269

van Driel, J., Gunseli, E., Meeter, M., & Olivers, C.N.L. (2017). Local and interregional alpha EEG dynamics dissociate between memory for search and memory for recognition. Neuroimage, 149, 114–128. https://doi.org/10.1016/j.neuroimage.2017.01.031

Varela, F., Lachaux, J.P., Rodriguez, E., & Martinerie, J. (2001). The brainweb: phase synchronization and large-scale integration. Nature Reviews. Neuroscience, 2, 229–239. https://doi.org/10.1038/35067550

Velo, J.R., Stewart, J.L., Hasler, B.P., Towers, D.N., & Allen, J.J. (2012). Should it matter when we record? Time of year and time of day as factors influencing frontal EEG asymmetry. Biological Psychology, 91, 283–291. https://doi.org/10.1016/j.biopsycho.2012.06.010

Vidaurre, D., Hunt, L.T., Quinn, A.J., Hunt, B.A.E., Brookes, M.J., Nobre, A.C., & Woolrich, M.W. (2018). Spontaneous cortical activity transiently organises into frequency specific phase-coupling networks. Nat Commun, 9, 2987. https://doi.org/10.1038/s41467-018-05316-z

Vinck, M., Oostenveld, R., van Wingerden, M., Battaglia, F., & Pennartz, C.M. (2011). An improved index of phase-synchronization for electrophysiological data in the presence of volume-conduction, noise and sample-size bias. Neuroimage, 55, 1548–1565. https://doi.org/10.1016/j.neuroimage.2011.01.055

Walter, D.O. (1963). Spectral analysis for electroencephalograms: mathematical determination of neurophysiological relationships from records of limited duration. Experimental Neurology, 8, 155–181. https://doi.org/10.1016/0014-4886(63)90042-6

Weisz, N., Hartmann, T., Muller, N., Lorenz, I., & Obleser, J. (2011). Alpha rhythms in audition: cognitive and clinical perspectives. Front Psychol, 2, 73. https://doi.org/10.3389/fpsyg.2011.00073

Wendling, F., Ansari-Asl, K., Bartolomei, F., & Senhadji, L. (2009). From EEG signals to brain connectivity: a model-based evaluation of interdependence measures. Journal of Neuroscience Methods, 183, 9–18. https://doi.org/10.1016/j.jneumeth.2009.04.021

Whitton, A.E., Deccy, S., Ironside, M.L., Kumar, P., Beltzer, M., & Pizzagalli, D.A. (2018). Electroencephalography Source Functional Connectivity Reveals Abnormal High-Frequency Communication Among Large-Scale Functional Networks in Depression. Biol Psychiatry Cogn Neurosci Neuroimaging, 3, 50–58. https://doi.org/10.1016/j.bpsc.2017.07.001

Whitton, A.E., Webb, C.A., Dillon, D.G., Kayser, J., Rutherford, A., Goer, F., Fava, M., McGrath, P., Weissman, M., Parsey, R., Adams, P., Trombello, J.M., Cooper, C., Deldin, P., Oquendo, M.A., McInnis, M.G., Carmody, T., Bruder, G., Trivedi, M.H., & Pizzagalli, D.A. (2019). Pretreatment Rostral Anterior Cingulate Cortex Connectivity With Salience Network Predicts Depression Recovery: Findings From the EMBARC Randomized Clinical Trial. Biological Psychiatry, 85, 872–880. https://doi.org/10.1016/j.biopsych.2018.12.007

Williams, L.M. (2017). Defining biotypes for depression and anxiety based on large-scale circuit dysfunction: a theoretical review of the evidence and future directions for clinical translation. Depression and Anxiety, 34, 9–24. https://doi.org/10.1002/da.22556

Womelsdorf, T., Schoffelen, J.M., Oostenveld, R., Singer, W., Desimone, R., Engel, A.K., & Fries, P. (2007). Modulation of neuronal interactions through neuronal synchronization. Science, 316, 1609–1612. https://doi.org/10.1126/science.1139597

Yeo, B.T., Krienen, F.M., Sepulcre, J., Sabuncu, M.R., Lashkari, D., Hollinshead, M., Roffman, J.L., Smoller, J.W., Zöllei, L., Polimeni, J.R., Fischl, B., Liu, H., & Buckner, R.L. (2011). The organization of the human cerebral cortex estimated by intrinsic functional connectivity. Journal of Neurophysiology, 106, 1125–1165. https://doi.org/10.1152/jn.00338.2011

Yokosawa, K., Murakami, Y., & Sato, H. (2020). Appearance and modulation of a reactive temporal-lobe 8-10-Hz tau-rhythm. Neuroscience Research, 150, 44–50. https://doi.org/10.1016/j.neures.2019.02.002

